# Haplotype-resolved genome and population genomics of the threatened garden dormouse in Europe

**DOI:** 10.1101/2024.02.21.581346

**Authors:** Paige A. Byerly, Alina von Thaden, Evgeny Leushkin, Leon Hilgers, Shenglin Liu, Sven Winter, Tilman Schell, Charlotte Gerheim, Alexander Ben Hamadou, Carola Greve, Christian Betz, Hanno J. Bolz, Sven Büchner, Johannes Lang, Holger Meinig, Eva Marie Famira-Parcsetich, Sarah P. Stubbe, Alice Mouton, Sandro Bertolino, Goedele Verbeylen, Thomas Briner, Lídia Freixas, Lorenzo Vinciguerra, Sarah A. Mueller, Carsten Nowak, Michael Hiller

**Author notes:** contributed equally to the manuscript (Paige A. Byerly, Alina von Thaden).

## Abstract

Genomic resources are important for evaluating genetic diversity and supporting conservation efforts. The garden dormouse (*Eliomys quercinus*) is a small rodent that has experienced one of the most severe modern population declines in Europe. We present a high-quality haplotype-resolved reference genome for the garden dormouse, and combine comprehensive short and long-read transcriptomics datasets with homology-based methods to generate a highly complete gene annotation. Demographic history analysis of the genome revealed a sharp population decline since the last interglacial, indicating an association between colder climates and population declines prior to anthropogenic influence. Using our genome and genetic data from 100 individuals, largely sampled in a citizen-science project across the contemporary range, we conducted the first population genomic analysis for this species. We found clear evidence for population structure across the species’ core Central European range. Notably, our data shows that the Alpine population, characterized by strong differentiation and reduced genetic diversity, is reproductively isolated from other regions and likely represents a differentiated evolutionary significant unit (ESU). The predominantly declining Eastern European populations also show signs of recent isolation, a pattern consistent with a range expansion from Western to Eastern Europe during the Holocene, leaving relict populations now facing local extinction. Overall, our findings suggest that garden dormouse conservation may be enhanced in Europe through designation of ESUs.

## Introduction

As genome sequencing technology advances and costs decrease, high-quality reference genomes are becoming increasingly recognized as an essential resource for conservation genomics (Formenti et al. 2022; Brandies et al. 2019; Theissinger et al. 2023; Paez et al. 2022). When aligned to a reference genome assembly, informative markers such as genome-wide single nucleotide polymorphisms (SNPs) generated via high-throughput genomic data can contribute to the understanding of past and contemporary population demographics (Gautier et al. 2016; du Plessis et al. 2023; Luo et al. 2023; Campana et al. 2016), reveal evolutionary patterns such as adaptive differentiation (Szarmach et al. 2021; Martchenko and Shafer 2023), and inform the current status of wildlife species of conservation concern (Talla et al. 2023; Viluma et al. 2022). While reduced-representation sequencing (RRS) methodologies continue to be the more cost-effective and efficient means of generating SNP datasets for large numbers of samples (Wright et al. 2020; Peterson et al. 2012), alignment of RRS reads to a high-quality reference genome improves both the precision of SNP calls and the quantity of SNPs recovered when compared to *de novo* read alignment without a genome assembly (Rochette et al. 2019; Brandies et al. 2019). Reference genomes are also useful for improving data recovery from closely-related species, with cross-species alignment shown to be highly effective (Takach et al. 2023; DeSaix et al. 2019; Burri et al. 2015; Nieto-Blázquez et al. 2022). Until recently, reference genomes for non-model organisms were often generated using short-read sequencing. However, short-read reference genomes are prone to errors such as sequence gaps, false duplicates, and misassembled reads, all of which can lead to inaccuracies in downstream data processing (Rhie et al. 2021). By contrast, genome assemblies generated from long-read sequencing are largely free from these sources of error. This higher accuracy combined with greater contiguity makes long-read sequencing more suitable for genomic analyses such as linking phenotype to genotype, detection of large structural variants, and gene expression analyses (Jebb et al. 2020; Rhie et al. 2021; Karimi et al. 2022; Wold et al. 2021). Given the value of high-quality reference genome assemblies for population and conservation genomic analyses, long-read sequencing of reference genomes for wildlife groups should be a focus of ongoing conservation efforts.

Highly-informative data sources such as reference genomes derived from long-read sequencing can have particular value for investigating understudied wildlife groups, as genomic data can contribute to baseline knowledge needed for conservation planning such as population size estimates and historical changes in demography. Within mammals, such foundational knowledge is lacking for many small-bodied taxa such as Rodentia and Eulipotyphla, which have a known deficit of research (Verde Arregoitia 2016; Kennerley et al. 2021) even in well-studied regions such as Central Europe (Pérez-Espona 2017). Indeed, general population trends of small mammals are not well understood in Europe (Lang et al. 2022; Pérez-Espona 2017; Gippoliti and Amori 2007; Bertolino et al. 2015), though many species appear to be declining (Rammou et al. 2022; Gippoliti and Amori 2007; Lang et al. 2022; Reiners et al. 2014). Population genomic analysis could expand our understanding of population structure and the factors contributing to declines in Europe, but with the exception of a few flagship species such as the Eurasian beaver (*Castor fiber*; (Halley et al. 2021)) it remains a limited tool in European rodent conservation.

The garden dormouse (*Eliomys quercinus*) is a small rodent species that exemplifies the small mammal conservation crisis in Europe. Once distributed across the continent, the garden dormouse is now recognized to have experienced one of the most extensive modern population declines on the European continent, with an approximate 51% range contraction since the 1970s (Bertolino 2017). Currently, it is only considered common in five of the 26 countries that once comprised its historical range, and even within these refuge countries its distribution is patchy and highly localized (Bertolino 2017). Despite its known imperiled status, research on the garden dormouse has been limited, even in comparison to the three other European species in its family Gliridae (Lang et al. 2022). In particular, the causes of its population decline are still unclear. Changing climate and intensification of land use have been suggested as possible influences; however, the demographic shift appears to have started prior to major 20th century landscape-level changes (Meinig and Büchner 2012; Bertolino 2017). Despite this possible influence of habitat loss, the garden dormouse also shows signs of being a habitat generalist, and this forest species has also adapted well to urban areas in some parts of its range while essentially vanishing from others (Meinig and Büchner 2012; Büchner et al. 2024), making its decline an ongoing mystery.

Population genetic analysis has been identified as a major research need in garden dormouse conservation (Meinig and Büchner 2012; Büchner et al. 2024), as genetic data could aid in understanding patterns of connectivity and gene flow between regions and uncover past population processes that may help explain current declines. Prior karyotyping and mitochondrial DNA analyses suggested the existence of four genetically distinct clades within Europe (Perez et al. 2013; Libois et al. 2012), although the existence of hybrid individuals indicated gene flow between clades (Perez et al. 2013). A better understanding of the population structure and genetic diversity within the hypothesized clades could help guide future management priorities and inform both regional and European-wide conservation efforts. Here, we present a high-quality long-read-based reference genome for the garden dormouse, representing the first such complete genome for both the species and its genus. We then demonstrate the utility of the genome by conducting the first genome-wide analysis of population differentiation across the contemporary range of the garden dormouse.

## Results

### Haplotype-resolved genome assembly

To generate a reference-quality assembly for the garden dormouse, we collected tissue from a male individual sourced from Mainz, Germany in 2023. We used DNA from this individual to generate 124.4 Gb of PacBio High-Fidelity (HiFi) long-read data, with 117 Gbp in long-range Hi-C read pairs. Assuming a 2.5 Gb genome size, similar to other dormice (Zoonomia Consortium 2020; Böhne et al. 2023), the HiFi and Hi-C data have an estimated coverage of ∼50× and ∼47×, respectively. With these data, we obtained two haplotype-resolved genome assemblies (Fig. S1).

The contig N50 values of the two haplotypes are 51.7 and 45.4 Mb, respectively. This contiguity is at least 2.8-fold higher compared with other Sciuromorpha genome assemblies (Fig. 1A,B, Table S1). Scaffold N50 values are 108.1 and 107.5 Mb. Haplotype 1 contains the X Chromosome, which was assembled as a chromosome-level scaffold. The Y Chromosome could not be assembled well. We identified 24 autosomes, implying that the sequenced individual has a karyotype of 50 chromosomes (2*×*24 + 2 sex chromosomes) (Fig. 1C,D), which matches the range of 48 to 54 chromosomes identified for *Eliomys* so far and corresponds to the expected karyotype of the Northern clade where the specimen was sampled (Perez et al. 2013). Overall, 99.4 and 99.6% of haplotype 1 and 2 are contained in chromosome-level scaffolds. These metrics exceed the standards set by the Vertebrate Genome Project (Rhie et al. 2021).

**Figure 1:**
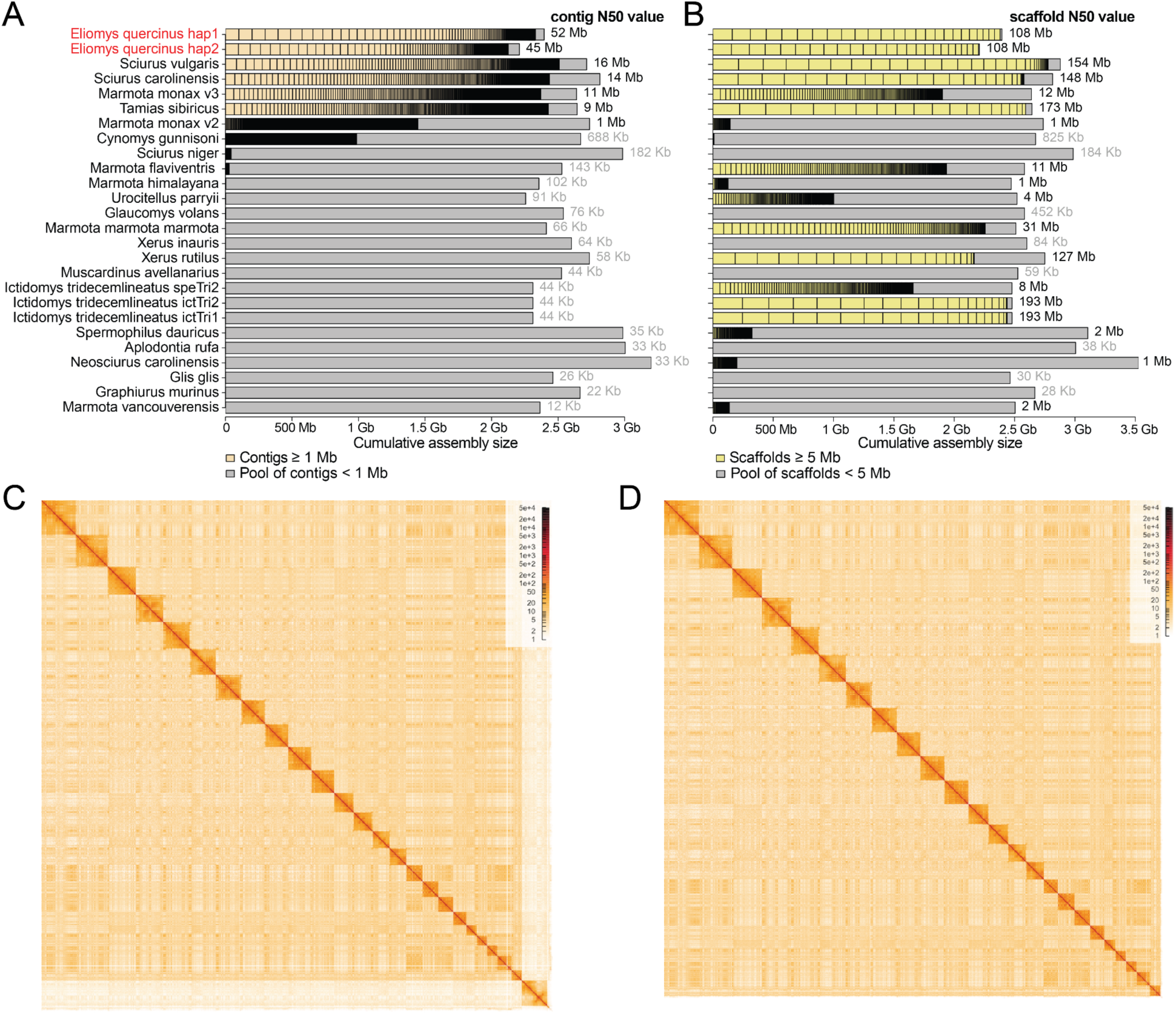
Contiguity and Hi-C maps of both haplotype assemblies. (A, B) Visualization of contig (A) and scaffold (B) sizes of the garden dormouse haplotype 1 and 2 (red font) and other Sciuromorpha genome assemblies. The N50 values are given on the right side. Contigs shorter than 1 Mb and scaffolds shorter than 5 Mb are not visualized individually, but shown as the gray portion of each bar. (C, D) Hi-C density maps for haplotype 1 (C) and haplotype 2 (D).

### Assembly quality and completeness

Using the HiFi reads, we estimate that our haplotype 1 and 2 assemblies have a high base accuracy with a QV (consensus quality) value of 60.46 and 60.24, indicating only one error per megabase. To assess gene completeness, we first used compleasm (Huang and Li 2023) with the BUSCO odb10 set of 9,226 near-universally conserved mammalian genes. Counting completely detected genes present in a single copy, our assemblies contained 98.3% such genes for haplotype 1 and 90.16% for haplotype 2 that lacks the X Chromosome. In comparison to other Sciuromorpha genome assemblies, our garden dormouse haplotype 1 ranks second after the *Sciurus carolinensis* assembly (Mead et al. 2020) (Fig. 2A, Table S1). Second, we used TOGA (Kirilenko et al. 2023) to compare the status of 18,430 ancestral placental mammal coding genes across the garden dormouse and other Sciuromorpha assemblies. TOGA explicitly distinguishes between the two major assembly issues (incompleteness and base errors) by classifying genes into those that have an intact reading frame, those that have gene-inactivating mutations (frameshifts, stop codons, splice site mutations, exon or gene deletions) and those that have missing sequences due to assembly incompleteness or fragmentation. We found that 94.1% of the ancestral genes are intact and only 1.1% have missing exons (Fig. 2B, Table S1). In comparison to other assemblies in the order Sciuromorpha, our garden dormouse haplotype 1 ranks third in terms of intact genes. Despite being top-ranked in the compleasm metric, TOGA reveals that the *S. carolinensis* assembly has substantially more genes with inactivating mutations, indicating a lower base accuracy (Fig. 2B).

**Figure 2:**
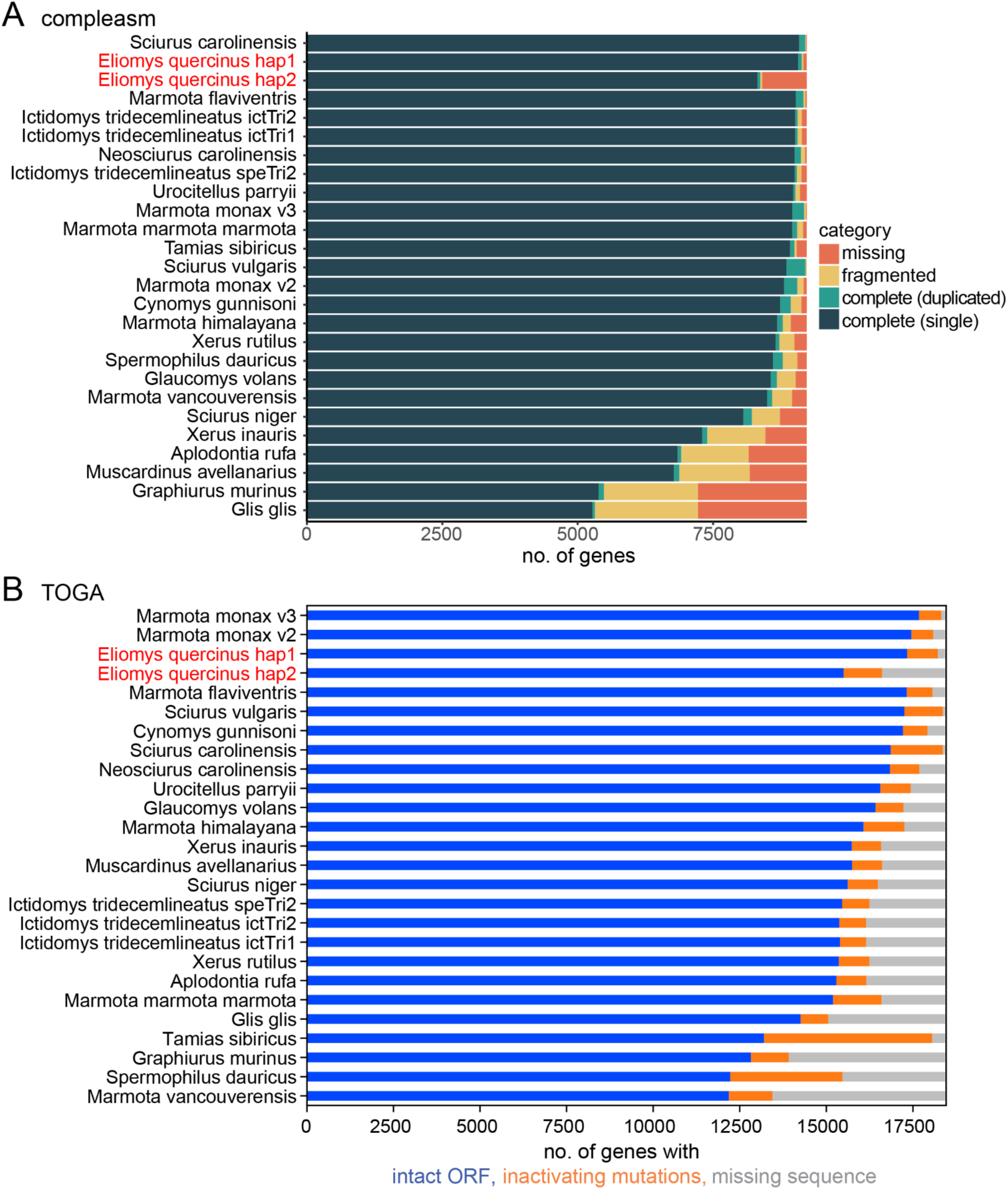
Comparison of gene completeness between our haplotype-resolved garden dormouse and other Sciuromorpha genomes. (A) Fractions of detected, fragmented, and missing BUSCO genes (mammalian odb10 dataset, 9,226 genes), as classified by compleasm v.0.2.2 (Huang and Li 2023) across Sciuromorpha assemblies. (B) TOGA classification of 18,430 ancestral placental mammal genes across Sciuromorpha assemblies. Assemblies are ranked by the number of completely-detected (A) and by the number of intact (B) genes. For the garden dormouse, we show both haplotypes in red font and next to each other, but the X Chromosome is only contained in haplotype 1. Species and assembly accessions are listed in Table S1.

### Gene annotation

To comprehensively annotate genes of the garden dormouse genome, we generated transcriptomics data from eight different organs (heart, testis, liver, lung, spleen, muscle, kidney, and bladder). We combined ∼4.2 billion short RNA-seq and ∼1.6 million long Iso-Seq reads to produce transcript models. Transcript models were joined with a comprehensive homology-based annotation, obtained by first generating genome alignments and then applying TOGA with human and mouse as the reference. The resulting annotation comprised 19,374 coding genes and as estimated by compleasm in protein mode, completely contains 99.91% of the mammalian BUSCO genes, which indicates a very high completeness.

### Long-term demographic history of the garden dormouse

To gain insight into the demographic history of the garden dormouse and investigate potential effects of climatic changes in the Pleistocene, we inferred the effective population size (N_e_) using pairwise sequentially Markovian coalescence (PSMC) (Li and Durbin 2011). We used a generation time of 1.5 years and a mutation rate of 5.7*×*10^-9^ per generation (see Methods).

Our results indicate relatively high N_e_ (> 130,000) and consistent population growth from 800 kya towards the last interglacial (Fig. 3). Population sizes reached their maximum of > 200,000 individuals around the last interglacial. Subsequently, N_e_ gradually declined towards the last glacial maximum, reaching an estimated minimum N_e_ of less than 10,000 individuals around 10 kya. Together, this indicates that the garden dormouse thrived in warm periods, while a cooling climate with longer winters caused severe population declines. Consistent with this result, the Gliridae family thrived during earlier warm periods in the Miocene (23 to 5.3 Mya) (Nadachoswki and Daoud 1995).

**Figure 3:**
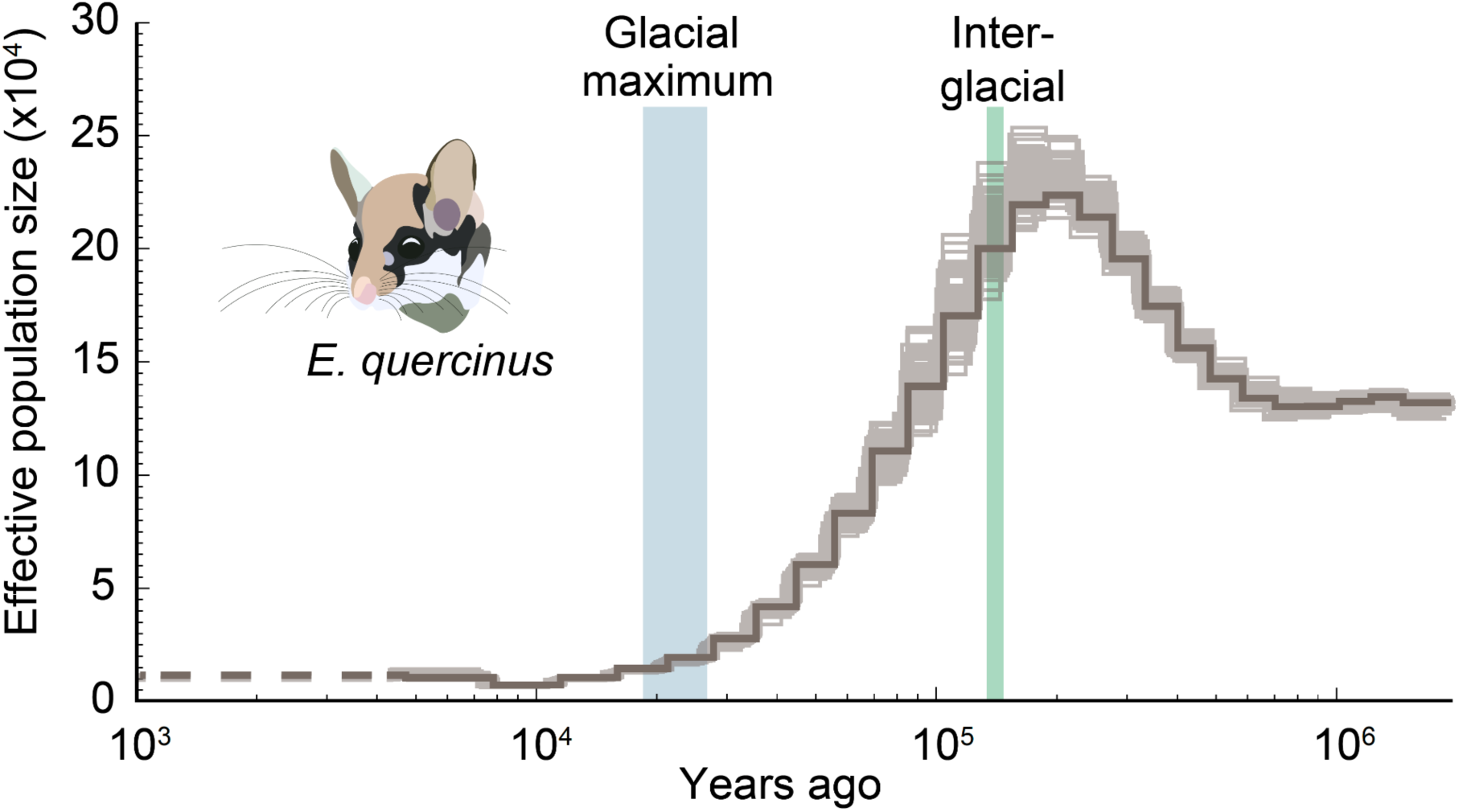
Demographic history of the garden dormouse. Effective population size (Ne) of *E. quercinus* as estimated with PSMC. N_e_ in ten thousands is shown on the y-axis and time in years ago on the x-axis. Faint lines indicate uncertainty of inferred N_e_ based on 100 bootstrap replicates. Blue and green bars show the timing of the last glacial maximum and last interglacial, respectively. The most recent N_e_ estimate with reduced accuracy is displayed by a dotted line on the left.

### Genotyping and variant calling for population genomics

To investigate population structure across the European range of the garden dormouse, we obtained *n* = 100 samples collected opportunistically between 1991–2020 for population genomic analyses and performed normalized Genotyping-by-Sequencing (nGBS). After sequencing, reads were aligned to the garden dormouse reference genome, and the resulting alignments were used to call single-nucleotide polymorphisms (SNPs). We assigned *a priori* population identity following in part the clades identified by (Perez et al. 2013) (Fig. 4A).

**Figure 4.**
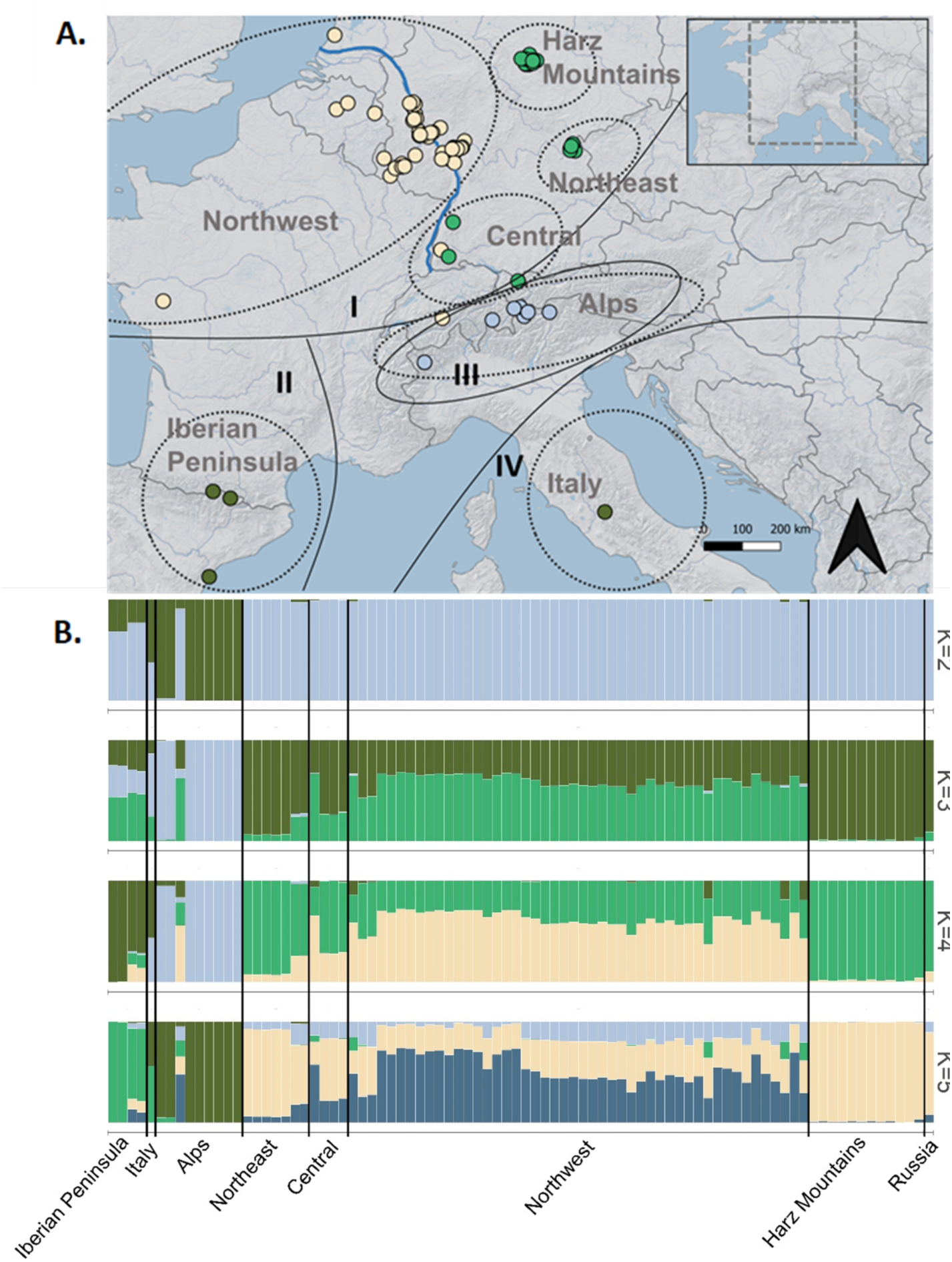
Garden dormouse population structure inferred from 47,115 nuclear SNP loci. (A) Group assignment via five PCs obtained by a discriminant analysis of principal components. The predicted assignment of genetic clusters (*K*) 1–3 is shown by four different colors and the *a priori* sampling region identity is highlighted by dotted gray lines and text. The clades I–IV that were previously identified based on mitochondrial DNA (Perez et al. 2013) are outlined in black. One sample from Russia is not pictured. The Rhine River is outlined in blue, and the boundary of the study area is outlined in gray in the inset map. (B) Genetic clustering inferred by Bayesian structure analysis for *K* ranging from 2 to 5 in STRUCTURE, with x-axes representing geographic sampling location and y-axes the proportion of group membership.

After removal of one individual with 98% missing data and one duplicate sample, we obtained data for a total of 41,175 SNPs from *n* = 98 samples with a mean per site depth of 14.75× and mean frequency of missingness of 0.11 (Table S2). For analyses dependent upon allele frequencies, such PCA-based analyses, we pruned SNPs based on linkage disequilibrium, resulting in a dataset of 19,022 SNPs.

We first inferred relatedness among individual samples and found six pairwise groups of first-order relations in the Harz population. Because inclusion of related individuals can bias population genomic analyses, we randomly removed one individual from each pair above a kinship threshold > 0.35, which corresponds to a first-order familial relationship, for subsequent analyses (Table S2). All Russian samples were identified as second order relations (kinship > 0.20), indicating half-sibling or grandparent-offspring relationships. Therefore we removed all but one individual from Russia, retaining the sample with the lowest missing data (Table S2). As all individuals with kinship greater than the threshold were collected on the same day from the same locations, these kinship estimates likely reflect accurate estimates of relatedness between individuals. This left us with a final dataset of *n* = 86 samples (Table S2). Because we had only one sample for Italy and Russia, we omitted these samples from further population genetic analyses, retaining these samples only for investigating population structure.

### Assessing genetic diversity

We quantified genetic diversity as both SNP-based (polymorphisms only) and genome-wide (both fixed and variable sites) heterozygosity and found highest genetic diversity for the Iberian Peninsula and Central regions (Table 1). Within the Central European region, genetic diversity was lower for population clusters east of the Rhine River (Northeast, Alps, Harz Mountains) than those located west of or near the river valley (Northwest, Central). This finding is consistent with the overall lower sizes of garden dormouse populations located east of the Rhine River in Central Europe (Bertolino 2017).

**Table 1.** Sample size (*n*) and genetic diversity metrics among eight garden dormouse sampling regions with number of SNP-based polymorphic sites, observed (H_obs_) and expected (H_exp_) heterozygosity, allelic richness rarified by sample size (A_R_), the inbreeding coefficient F_IS_, and nucleotide diversity *π* estimated from 19,022 SNP loci, and observed autosomal heterozygosity (H_obs_A_) estimated from from ∼90,412 variant and ∼58,406,409 fixed sites. Values for the Northwest region were generated by running four separate analyses of *n* = 12 and then averaging the output to avoid biases resulting from the uneven sample size between the Northwest (*n* = 48) and the other regions.

| Region | *n* | polymorphic sites | $H_{obs}$ | $H_{exp}$ | $F_{IS}$ | $\pi$ | $H_{obs\_A}$ |
| --- | --- | --- | --- | --- | --- | --- | --- |
| Alps | 9 | 5975 | 0.036 | 0.068 | 0.149 | 0.075 | 0.00028 |
| Central | 4 | 13552 | 0.257 | 0.273 | 0.123 | 0.321 | 0.00017 |
| Harz Mountains | 12 | 12992 | 0.223 | 0.233 | 0.053 | 0.245 | 0.00034 |
| Iberian Peninsula | 4 | 11389 | 0.249 | 0.230 | 0.056 | 0.280 | 0.00067 |
| Northeast | 7 | 13173 | 0.209 | 0.244 | 0.124 | 0.264 | 0.00036 |
| Northwest | 48 | 18062 | 0.286 | 0.344 | 0.180 | 0.360 | 0.00042 |

*F*_ST_ was significantly high (confidence intervals did not overlap 0) between most regional pairwise groupings (Table 2), suggesting that dispersal between spatially disjunct populations may be limited. However, the highest overall *F*_ST_ values were consistently recovered for pairwise comparisons of the Alpine group and all other populations.

**Table 2.** Pairwise *F*_ST_ between six garden dormouse sampling regions below the diagonal and 95% confidence intervals based on 1,000 bootstraps above the diagonal, corrected for sample size. Sampling regions are defined and abbreviated as Northwest (NW), the Alps (ALP), Harz Mountains (HAR), Central (CE), the Iberian Peninsula (IB), and Northeast (NE).

|  | NW | ALP | HARZ | CE | IB | NE |
| --- | --- | --- | --- | --- | --- | --- |
| NW | — | 0.336, 0.344 | 0.162, 0.168 | 0.063, 0.068 | 0.187, 0.195 | 0.133, 0.139 |
| ALP | 0.340 | — | 0.564, 0.576 | 0.556, 0.568 | 0.472, 0.488 | 0.560, 0.571 |
| HARZ | 0.165 | 0.570 | — | 0.208, 0.219 | 0.377, 0.389 | 0.154, 0.163 |
| CE | 0.065 | 0.562 | 0.213 | — | 0.245, 0.259 | 0.083, 0.093 |
| IB | 0.191 | 0.480 | 0.383 | 0.252 | — | 0.331, 0.344 |
| NE | 0.136 | 0.565 | 0.159 | 0.088 | 0.337 | — |

### Assessing population structure

Phylogenetic analysis using RAxML showed that individuals from Northeastern Central Europe (Harz Mountains, Central, Northeast, Russia) form a single clade, separated from the Northwest, Alps, Italy, and Iberian Peninsula regions (Fig. S2) except for a single Central individual (G190084BW), which clustered with the Northwest group. This tree conformed with the PCA of genetic distance among individuals and sampling regions. PCA showed a strong separation between the Alpine and other sampling regions, with this differentiation explaining 14.48% of the variation on PC1 (Fig. 5). Further spatial differentiation between the Northwest region and the other Central European sampling locations explained 8.19% of the variation on PC2. These results were consistent with findings of relatively high *F*_ST_ among regions.

**Figure 5.**
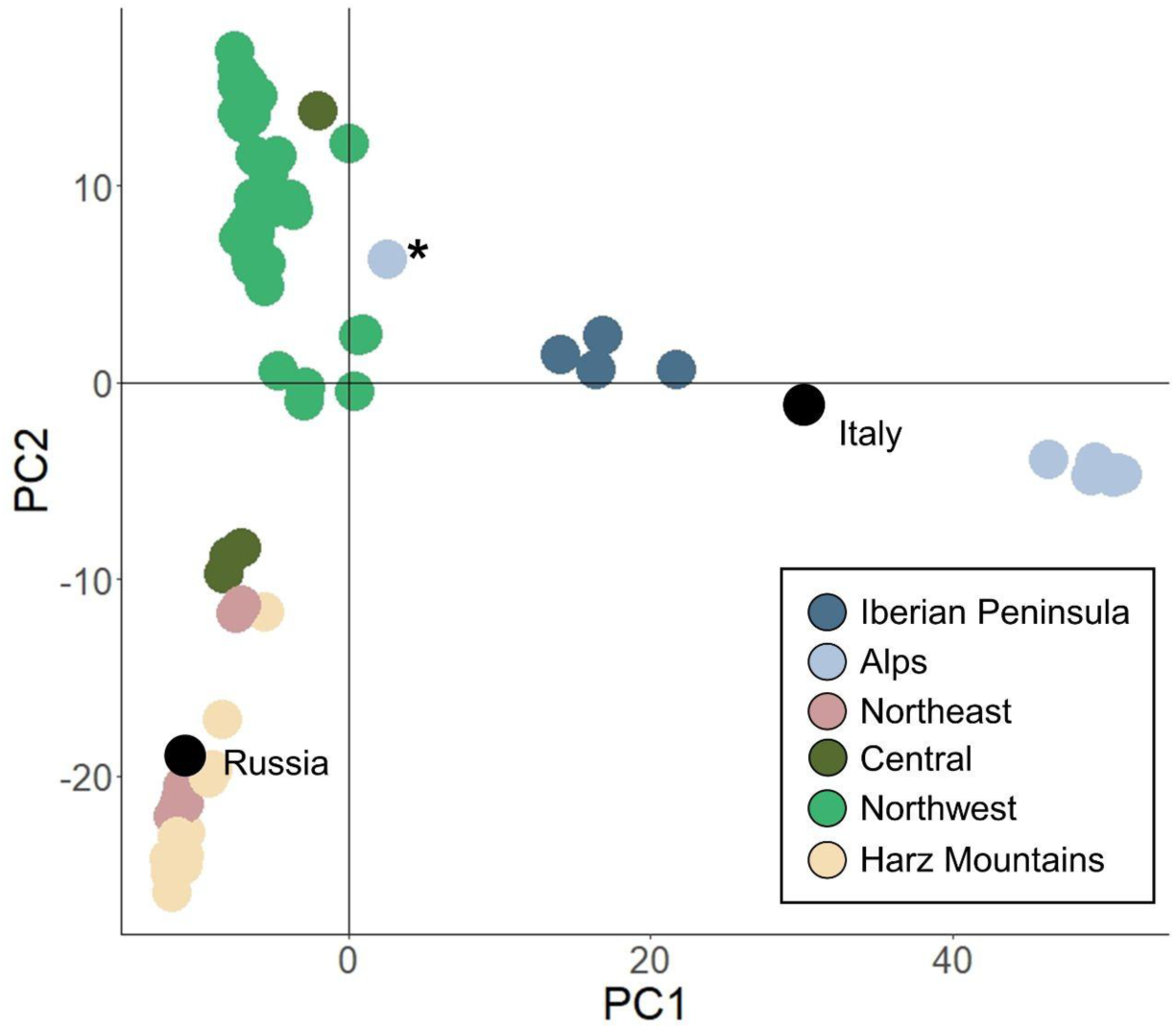
Principal component analysis (PCA) for the garden dormouse. The PCA is based on 19,022 nuclear SNP loci, with colors representing the geographical origin. Results are plotted on the first two principal components, with PC1 explaining 14.48% and PC2 8.19% of variation within the dataset. Regions where *n* = 1 (Italy and Russia) are indicated by black circles, with region names labeled within the plot. One individual from the Alps clustered separately from the Alpine region and is marked with a **\***.

We used DAPC and STRUCTURE to infer the number of genetic clusters within our samples. DAPC identified *K* = 4 as the most probable number of genetic clusters (Fig. S3A), while the results for STRUCTURE were less clear, suggesting *K* = 2–4 as possible cluster numbers (Fig. S4). Cluster membership for DAPC was suggestive of relatively strong differentiation between the Northwest region versus other regions in Germany (Fig. 4A, Fig. S3B,C), while STRUCTURE indicated greater admixture among these regions at all values of *K* (Fig. 4B; Fig. S5). Both methods were consistent in differentiating the Alpine samples from all other regions (Fig. 4B; Fig. S5). Patterns observed for the full dataset were comparable for the STRUCTURE analysis subset by sample size (Fig. S6), indicating that our results were not influenced by unequal sample size among our sampling regions for regions where n > 1.

Overall, population genomic analyses consistently revealed little evidence for contemporary gene flow between the Alpine region and other populations, and the high degree of differentiation in this region may indicate long-term isolation of the Alpine group. Despite the clear isolation of the Alpine group, all analyses were consistent in separating one individual from other samples collected from the Alpine region (G190088CH). This individual, which was collected north of the main Alpine ridge, instead showed strong evidence for admixture with the Northwest and Harz Mountains regions (Fig. 4; Fig. 5).

### Recent demographic history inferred from the site-frequency spectrum

To infer recent demographic histories and N_e_ for each individual population, we used coalescent simulations and compared three models (Fig. S7): constant steady population size (null model), population bottleneck (instantaneous size change model), and constant population growth or decline (incremental size change model). For all populations, the instantaneous size change (bottleneck) model showed the strongest support based on the lowest AIC value, although likelihood differences between models were not great (Table 3). The timing of the bottlenecks was largely consistent among the Alps, the Harz, and the Northeast, indicating a bottleneck event approximately ∼3,000 years ago. The Northwest model presented a slightly more distant bottleneck, at 7,365 years ago, which aligns with the population minimum ∼10,000 years ago inferred by PSMC. For the more northerly populations Harz, Northeast, and Northwest, differences between N_e_ before and during the bottleneck were not extensive (Table 3), indicating that the inferred population change may have been more gradual, which is in line with the steady decline suggested by PSMC. Despite this, all populations show contrastingly low contemporary N_e_, with the lowest N_e_ inferred for the Harz and Northeast populations. By contrast, the top model indicated that the Alps population experienced a catastrophic decline during the bottleneck, with N_e_ reduced by 99.99% before recovering to over 2x ancestral population size.

**Table 3.**
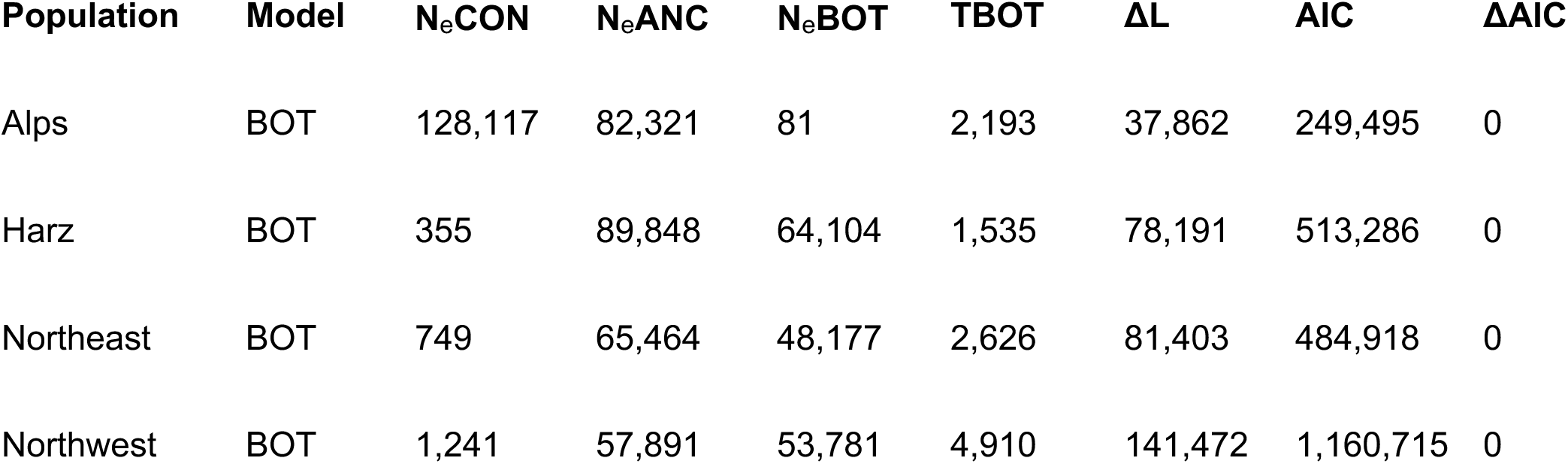
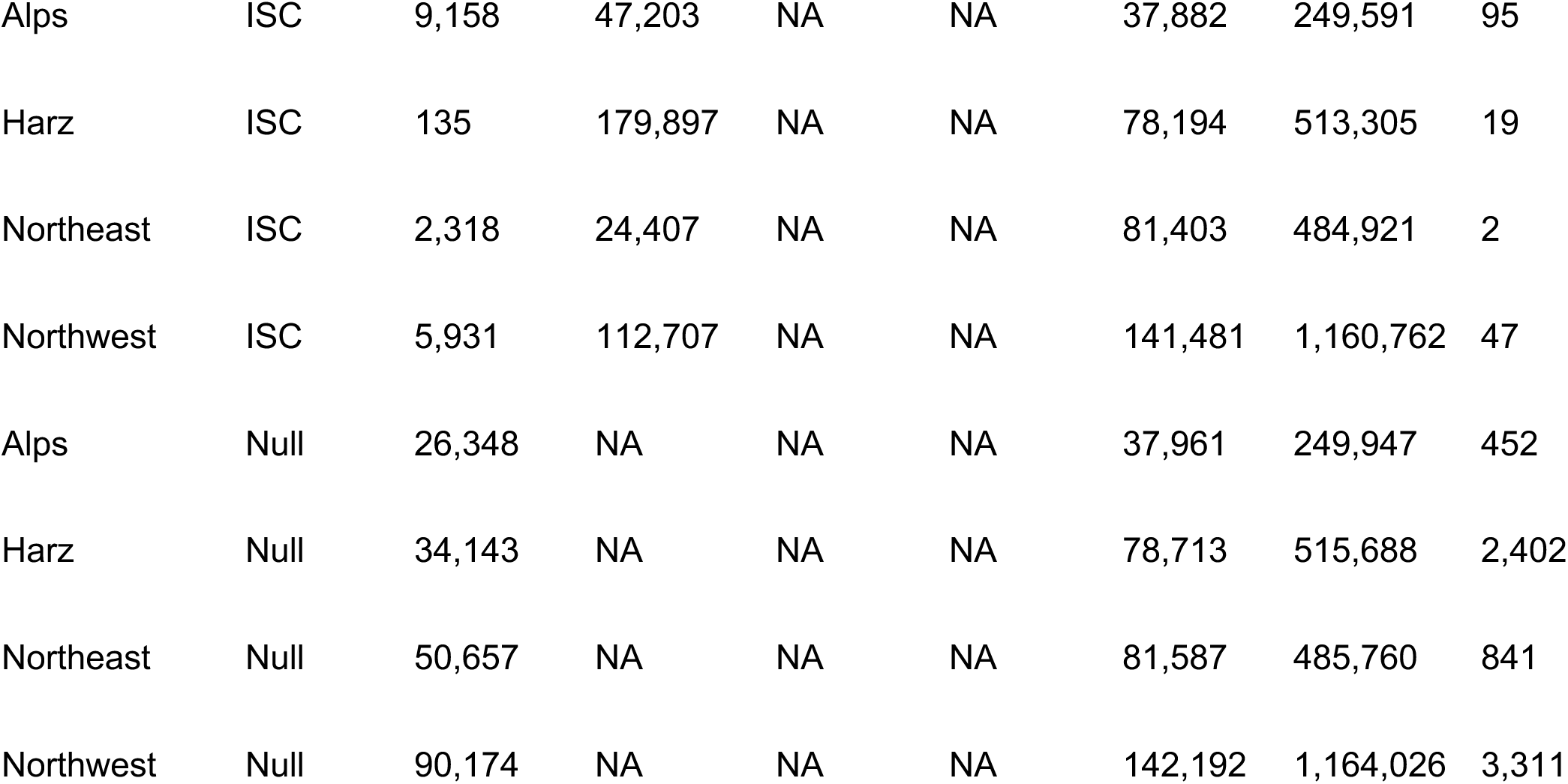
Model selection results from demographic inference using the Site Frequency Spectrum (SFS) in *fastsimcoal2*. Three models were run for each regional grouping of Alps, Harz, Northeast, and Northwest (Population). Model refers to one of three model types: Null model of constant population size (Null), Instantaneous Size Change/Bottleneck (BOT), and Incremental Size Change (ISC). Parameters were estimated for contemporary N_e_ (N_e_CON), ancestral N_e_ (N_e_ANC), N_e_ during bottleneck (N_e_BOT), and time of bottleneck in number of generations (TBOT). Model fit and selection criterion are shown as delta likelihood, calculated as maximum estimated likelihood maximum observed likelihood (ΔL), AIC, and ΔAIC.

| Population | Model | $N_e$CON | $N_e$ANC | $N_e$BOT | TBOT | $\Delta L$ | AIC | $\Delta AIC$ |
| --- | --- | --- | --- | --- | --- | --- | --- | --- |
| Alps | BOT | 128,117 | 82,321 | 81 | 2,193 | 37,862 | 249,495 | 0 |
| Harz | BOT | 355 | 89,848 | 64,104 | 1,535 | 78,191 | 513,286 | 0 |
| Northeast | BOT | 749 | 65,464 | 48,177 | 2,626 | 81,403 | 484,918 | 0 |
| Northwest | BOT | 1,241 | 57,891 | 53,781 | 4,910 | 141,472 | 1,160,715 | 0 |
| Alps | ISC | 9,158 | 47,203 | NA | NA | 37,882 | 249,591 | 95 |
| Harz | ISC | 135 | 179,897 | NA | NA | 78,194 | 513,305 | 19 |
| Northeast | ISC | 2,318 | 24,407 | NA | NA | 81,403 | 484,921 | 2 |
| Northwest | ISC | 5,931 | 112,707 | NA | NA | 141,481 | 1,160,762 | 47 |
| Alps | Null | 26,348 | NA | NA | NA | 37,961 | 249,947 | 452 |
| Harz | Null | 34,143 | NA | NA | NA | 78,713 | 515,688 | 2,402 |
| Northeast | Null | 50,657 | NA | NA | NA | 81,587 | 485,760 | 841 |
| Northwest | Null | 90,174 | NA | NA | NA | 142,192 | 1,164,026 | 3,311 |

## Discussion

By generating a long-read-based high-quality reference genome for the garden dormouse, our study provides a genetic resource that can be used to aid conservation efforts of small mammals in Europe. This study was performed in the frame of the citizen-science focused project *Spurensuche Gartenschläfer* (‘In Search of the Garden Dormouse’), which is the first national conservation and research project for this species in Germany (Büchner et al. 2024). The project aims to restore garden dormouse populations, as the species continues to decline across much of its range (Lang et al. 2022). Using our long-read genome assembly and population genomics data from samples across the species’ range, we were able to investigate garden dormouse demographic history, as well as contemporary population structure and genetic diversity parameters.

The long-term demographic history inferred with PSMC indicates that garden dormice underwent a substantial population bottleneck prior to modern anthropogenic influence, with a sustained decline and long-term low N_e_ following the last interglacial. Our data suggest that the warm climate around the last interglacial period supported high effective population sizes, and that hibernation did not save garden dormouse populations from declining during the last ice age. Previous paleontological data suggested that Gliridae diversified during periods of high glaciation (Lu et al. 2021). This observation has been interpreted as a sign that this family thrived during cold periods, likely because hibernation was an evolutionary advantage to survive cooling climates (Lu et al. 2021). In contrast, our results provide evidence that garden dormouse populations declined by more than 90% as the climate cooled down towards the last glacial maximum, raising the possibility that the observed lineage diversification during glaciation may have been more likely due to population fragmentation. This finding is in line with longer winters leading to reduced fitness following later emergence from hibernation (Lane et al. 2012). Similar results of declining populations after the last interglacial period have further been described for other Sciuromorpha, including the hibernating Gunnison’s prairie dog *Cynomys gunnisoni* (Tsuchiya et al. 2020) and the southern flying squirrel *Glaucomys volans* (Wolf et al. 2022)

More recent demographic history inferred with our SNP dataset further supports a bottleneck history for the individual populations analyzed. While the populations of Harz, Northwest, and Northeast showed evidence for moderate reductions of their ancestral population sizes during the historical declines, estimates of contemporary population size indicated a significant reduction. It is conceivable that the more recent demographic trajectories of these regions were of a complex nature that our models were not able to approximate; however, the low contemporary population sizes as compared to ancestral do reflect what is known about the extensive garden dormouse declines in the 20th and 21st centuries. By contrast, the Alps population showed signs of a massive and severe bottleneck with a more than 99.99% population reduction, followed by a more than two-fold modern recovery. While a severe bottleneck may in part explain the lower genetic diversity of this region, the magnitude of the decline seems too extreme and the estimated contemporary N_e_ inappropriately high for the Alpine region, which may indicate that none of the models tested accurately represented the demography of the Alpine population. While the accuracy and precision of *fastsimcoal2* is known to be affected by the number of SNPs analyzed and the amount of missing data (Nunziata and Weisrock 2018), *fastsimcoal2* is also sensitive to population census size, and may not accurately detect population declines when census size is low (Nunziata and Weisrock 2018). It is unclear how these and other factors, such as sample size and temporal heterogeneity, may influence model output (Marchi et al. 2024). The Alps sample set spans a collection period of 19 years, covers a wide geographic area, and exhibits extremely low genetic diversity. These factors may have influenced parameter output for this region and led to upwardly biased N_e_. Future analysis with whole genome resequencing is needed to better explore the demographic history of the species, and of the Alpine region in particular.

Our findings of high genetic differentiation across the garden dormouse’s core Central European range and the existence of four genetic clusters confirm and extend prior genetic research, which also identified four garden dormouse lineages within a similar sampling area based on mitochondrial DNA haplotypes and chromosome number differences (Perez et al. 2013; Forcina et al. 2022). As in Perez et al. 2013, we found the Alpine region south of the main Alpine ridge to be strongly differentiated from all other sampling regions, as evidenced in our study by both model-based (STRUCTURE), and non-model-based (PCA) approaches. This strong signal of genetic differentiation displayed by the Alpine population further confirms prior mtDNA genetic distance estimates and karyotypic divergence between the Alps and other regions across the garden dormouse’s range (2*n* = 52 “eastern Alps” and 54 “western Alps”) (Perez et al. 2013). One specimen collected within our Alps boundary showed admixture with more Northern regions, but this specimen was collected north of the main Alpine ridge and likely does not belong to the highly differentiated Alpine population. We additionally found the regions south of the Alpine range to be differentiated from those north of the range, which supports that this mountain range may act as a barrier to garden dormouse dispersal (Perez et al. 2013). In contrast to previous research, our more comprehensive datasets also allowed us to detect additional moderate population structure within the Central European clade (Fig. 4), showing a clear pattern of differentiation between eastern and western Central Europe.

Combined with previous findings of reciprocal mtDNA monophyly (Perez et al. 2013), the significant differentiation of nuclear loci found in this study supports the designation of the Alpine garden dormouse population as an evolutionary significant unit (ESU) according to the classic conservation genetics definition (Moritz 1994). Within the Alpine region, the garden dormouse is found in both subalpine and montane habitats, where it occupies coniferous as well as deciduous forest habitat (Bertolino 2017). The strong genetic differentiation and distinct habitat suggests the potential for long-term genetic separation, possibly arising from adaptation to local habitat conditions or climate. Future research should focus on investigating potential adaptive differences with whole genome resequencing. Although the garden dormouse has been largely reported as demographically stable in the Alps (Bertolino 2017), our findings of low genetic diversity and high rates of inbreeding are indicative of small population sizes, and may indicate that population declines in this area have been greater than previously thought. Hence, the status of this population should be reevaluated. Perez et al. (2013) also found lower nucleotide diversity for this region via mtDNA analyses, and surmised that reductions in genetic diversity may have resulted from population expansion following a genetic bottleneck, which our results also support. The proportional differences in genetic diversity parameters between the Alpine and other populations and other sampling regions reported in this study are comparable to those seen in comparisons between mainland and both oceanic and montane “habitat” island populations of other threatened small mammalian species (Rick et al. 2023; White and Searle 2007; Ditto and Frey 2007), providing further support for the Alpine population as an isolated, genetically distinct unit that may require the implementation of separate conservation strategies.

The range contraction of the garden dormouse is known to be centered in the eastern part of its range, with the species becoming more common and populations more contiguous as one moves further west (Bertolino 2017; Meinig and Büchner 2012). Fossil evidence indicates that the garden dormouse originated on the Iberian Peninsula and spread out into Central Europe during the Holocene (Mansino et al. 2014), which may mean that remaining garden dormice in Eastern Europe represent small, relict populations on the edge of extirpation (Anděra 1986). This is somewhat supported by the higher autosomal genetic diversity and lower inbreeding estimate we detected for the westernmost Iberian clade, which reflects a larger and more connected population. However, while the garden dormouse is still regarded as relatively common in some parts of the Iberian Peninsula (Bertolino 2017), extensive declines and local extirpations have been noted in some parts of Portugal and Spain, and the distribution has been characterized as patchy and scattered across the region (Santoro et al. 2017; Vale-Gonçalves and Mira 2023; Pita et al. 2021). A comparative analysis using the entire mitogenome found that the Iberian clade exhibits relatively lower genetic diversity as compared to that of Central Europe, with the most genetically diverse individuals instead centered in France, possibly resulting from admixture following emergence from glacial refugia in central Europe (Forcina et al. 2022). Given that samples for our study were collected from the northernmost boundary of the Iberian Peninsula, our characterization of the Iberian clade may therefore be more reflective of this contiguous core population rather than of the wider peninsula. This sampling distribution likely also explains the minimal genetic structuring we found between the Iberian Peninsula and the central European regions, which contrasts with the strong differentiation shown in mitochondrial studies with broader sampling distributions (Forcina et al. 2022; Perez et al. 2013).

Given the known contractions and declines in the eastern edge of the garden dormouse’s range, it is likely that the lower genetic diversity of the eastern populations in our study is reflective of genetic drift resulting from modern isolation of these populations. Support for this is provided by our observation of reduced genetic diversity and greater rates of inbreeding for the more easterly populations in Germany (Harz mountains and Northeast), whereas the Central population shows evidence of admixture with the larger and more contiguous Northwest population, thereby potentially increasing its genetic diversity through interbreeding. Larger sampling efforts are required to substantiate these preliminary findings. Notably, a similar east-west pattern of differentiation has also been observed for the hazel dormouse (*Muscardinus avellanarius*), which overlaps the garden dormouse range and is distributed as two separate lineages or ‘partial ESUs’, delineated along a similar geographical divide to the garden dormouse (Leyhausen et al. 2022; Mouton et al. 2017). Unlike the hazel dormouse, however, this east-west pattern of differentiation was not reflected by mtDNA in the garden dormouse (Perez et al. 2013), indicating that the division among lineages may be a more recent phenomenon resulting from habitat fragmentation and/or demographic changes such as population declines.

Although the causes of the garden dormouse’s east-to-west pattern of range contraction are not fully understood, landscape level changes driven by land use and climate change have been partly implicated (Bertolino 2017). However, it is unclear what specific role climate might play in this contraction. For the ecologically-similar hazel dormouse, warmer and wetter winters have been found to have a negative effect on adult survivorship, possibly because warmer temperatures cause animals to wake during hibernation, thereby expending energy and reducing fat reserves (Combe et al. 2023). Garden dormice hibernate in regions of harsh winter, and their life histories appear strongly tied to winter temperatures, as well as the onset and duration of the season (Bennett and Richard 2021; Mahlert et al. 2018). Timing and duration of activity periods and hibernation vary widely based on local climate conditions, with Mediterranean populations hibernating only 1–2 months per year, if at all (Viñals-Domingo et al. 2020), while Alpine populations may hibernate for 7 months per year (Bertolino et al. 2001). Further investigation of the role of climate in shaping the garden dormouse’s population structure via local adaptation may be useful in predicting the future viability of the species with ongoing climate change.

The garden dormouse is currently listed as “Near Threatened” in the International Union for Conservation of Nature (IUCN) Red List (Bertolino et al. 2008), but its status will be changed to “Vulnerable”(Bertolino 2017)in the next assessment (Bertolino 2017) due to the observed continuous decline in the species’ range. Our results support this change in status, and suggest that the garden dormouse in Europe is threatened due to population isolation and low regional genetic diversity, especially at the eastern edge of its range. Hence, intervention is needed to mitigate further population extirpations. We further provide strong evidence for the designation of the Alpine subpopulation of the garden dormouse as a separate ESU. Further research is needed to determine if local adaptation is driving the genetic differentiation of this and other isolated garden dormouse populations, and how these potential adaptations may influence future population viability in the face of a changing climate. Our annotated genome will enable comparative analyses addressing open evolutionary questions of the *Eliomys* species complex, as well as inform specific conservation planning and management for the garden dormouse in Europe. We hope that the findings outlined in this study can help guide future research for the garden dormouse, while highlighting the need for further protection within regions of population isolation and/or low genetic diversity.

## Materials and Methods

### Sample collection

Tissue samples for nGBS were collected from across the European distribution in the frame of a larger conservation project on *Eliomys quercinus* in Germany. Samples originated from Germany (*n* = 66), other Central European countries (*n* = 25), and Russia (*n* = 7) and were collected between 1991-2020 (see Table S2). All samples were taken opportunistically from dead-found individuals brought to museums or animal shelters, during monitoring, or during citizen-science activities, and all in compliance with respective local and national laws.

### Genome sequencing

To sequence a high-quality genome, we used a male individual collected in March 2023 in Mainz that was euthanized by a veterinarian due to injuries. A voucher specimen for this individual was prepared and is housed at the Zoological Research Museum Alexander Koenig under catalog number ZFMK MAM 2024.0263. DNA extraction, PacBio and Hi-C sequencing followed standard protocols that are described in the Supplementary Methods.

### Transcriptome sequencing

We generated RNA-and Iso-Seq transcriptomics data from the same individual, following standard protocols that are described in the Supplementary Methods.

### Genotyping-by-Sequencing

Tissue samples for normalized genotyping-by-sequencing (nGBS) were extracted using the DNeasy Blood & Tissue Kit (Qiagen) with slight protocol modifications to increase yield of RNA-free genomic DNA as described in (Leyhausen et al. 2022). Extracts were quantified with fluorometry (Qubit) and spectrophotometry (NanoDrop) and screened for DNA integrity using gel electrophoresis to identify samples of sufficient quality for high-throughput sequencing. Selected samples (*n* = 100) with ≥ 300 ng of high molecular weight DNA were sent for nGBS (Restriction Associated DNA sequencing [RAD-Seq]) at LGC Genomics GmbH (www.biosearchtech.com). Briefly, nGBS methodologies included enzymatic digestion with the enzyme MslI; purification; barcoding, library preparation, and PCR amplification with Illumina TruSeq amplification primers; size selection for 300–500 bp with the BluePippin system; and sequencing on an Illumina NextSeq 500/550. Sequences were then demultiplexed in house by LGC Genomics using proprietary software. Further details on enzymatic digestion, library preparation and nGBS protocols can be found in (Leyhausen et al. 2022; Arvidsson et al. 2016).

### Genome assembly

We produced haplotype-specific assemblies from HiFi reads using hifiasm (v.0.19.5), integrating Hi-C reads for phasing (Cheng et al. 2021). Both haplotypes were subject to filtering procedure using Foreign Contamination Screen Tool (FCS) (Astashyn et al. 2024). Contamination free assemblies were then polished to remove unambiguous heterozygous sites. Specifically, we mapped HiFi reads back to each assembly with minimap2 v.2.26 (Li 2018), removed duplicates using Picard MarkDuplicates tool v.3.1.0 (http://broadinstitute.github.io/picard), and called variants using DeepVariant v1.5.0 (Poplin et al. 2018). We then filtered for sites with genotype 1/1 and a ’PASS’ filter value, meaning that all or nearly all reads support an alternative sequence at this position and passed DeepVariants internal filters. Finally, we corrected corresponding nucleotide sites in the assembly using BCFtools consensus v.1.13 (Li 2011).

To scaffold the polished haplotype assemblies, we first mapped the Arima Hi-C reads to each of the two assemblies independently using chromap v.0.2.5 (Zhang et al. 2021) and then processed mapped reads with yahs v1.1a (Zhou et al. 2023). Finally, we performed manual curation of the two scaffolded haplotypes jointly analyzing them in PretextView v.0.2.5 (https://github.com/wtsi-hpag/PretextView). To this end, we remapped the Hi-C reads to both haplotype assemblies concatenated together using again chromap but allowing for multi-mapping reads (-q 0) to avoid discarding information in regions identical between two haplotypes. We also identified telomeric sequences with tidk v.0.2.31 (https://github.com/tolkit/telomeric-identifier) and where necessary corrected wrong contigs orientations to have telomeres in the ends of resulting scaffolds. To finalize changes made via manual dual haplotype curation, as recently proposed by the Vertebrate Genome and Darwin Tree of Life Project, we used the rapid curation framework (https://gitlab.com/wtsi-grit/rapid-curation/-/tree/main) from the Genome Reference Informatics Team (Howe et al. 2021).

To estimate the assembly base accuracy, we used Merqury (Rhie et al. 2020) with the HiFi reads. It should be noted that the QV estimates represent an upper bound as they are based on the HiFi reads used for assembly. To assess gene completeness, we used compleasm 0.2.2 (Huang and Li 2023) with the BUSCO odb10 set of 9,226 near-universally conserved mammalian genes and TOGA (below).

### Repeat masking and whole-genome alignment

RepeatModeler version 2.0.4 (default parameters) was used to generate a *de novo* repeat library for the garden dormouse assembly. RepeatMasker 4.1.0 (parameters - engine ncbi) was used with the resulting library to soft-mask the genome.

We followed our previous workflow to align the human (hg38 assembly) and the mouse (mm10) assembly to both garden dormouse haplotype assemblies (Sharma and Hiller 2017; Blumer et al. 2022). Briefly, we used LASTZ version 1.04.15 with sensitive parameters (K = 2400, L = 3000, Y = 9400, H = 2000, and the LASTZ default scoring matrix) (Harris 2007), axtChain (default parameters except linear-Gap=loose) to compute co-linear alignment chains (Kent et al. 2003), RepeatFiller (default parameters) to capture additional alignments between repetitive regions (Osipova et al. 2019), and chainCleaner (default parameters except minBrokenChainScore = 75,000 and -doPairs) to improve alignment specificity (Suarez et al. 2017).

### Gene annotation

We used TOGA version 1.1.4 (Kirilenko et al. 2023) with the alignment chains and the human GENCODE 38 and the mouse GENCODE M25 annotation (Frankish et al. 2021) to annotate coding genes. We ran TOGA for both garden dormouse haplotypes.

Raw RNA-seq data were processed with fastp v0.23.4 (Chen 2023; Chen et al. 2018) to remove adapters and low quality (< Q15) bases. Processed reads were mapped to a first haplotype assembly with HISAT2 v.2.2.1 (Kim et al. 2019). For Iso-Seq reads a circular consensus was generated from the subreads using ccs v.6.4.0 (https://github.com/PacificBiosciences/pbbioconda). Adapter trimming and base quality filtering was performed with lima v2.2.0 (https://github.com/PacificBiosciences/pbbioconda). Processed reads were cleaned from polyA tails and polymerase switching artifacts using ‘refine’ command from the Iso-Seq package v.4.0.0 (https://github.com/PacificBiosciences/IsoSeq/blob/master/isoseq-clustering.md) and then clustered together and aligned to generate consensus reads using the ‘cluster’ command from the Iso-Seq package. Additionally, reads were filtered for alignment quality using a custom Perl script (Supplementary Code). To combine transcriptome information from RNA-seq and Iso-Seq, we then used a hybrid option (--mix) in StringTie v.2.1.2 (Shumate et al. 2022).

To integrate transcriptomics and homology-based transcript evidence, we first merged human and mouse TOGA, keeping only transcripts with an intact reading frame.

We then added transcripts classified as partially intact and uncertain loss, but only if they do not overlap intact transcripts. The RNA- and Iso-Seq merged transcriptome data was then used to add UTRs to TOGA-annotated coding transcripts with a compatible exon-intron structure. Finally, we added transcriptome data for genomic loci that don’t have a TOGA transcript prediction to incorporate lineage-specific and non-coding transcripts into the annotation.

### Demographic history inferred from the reference genome

We used PSMC (Li and Durbin 2011) to infer the demographic history of the garden dormouse across the past one million years. Minimap2 v.2.26 (Li 2018) was used to map the PacBio HiFi reads to haplotype 1. Heterozygous sites were called with DeepVariant v1.2.0 (Poplin et al. 2018) using the PacBio model. A consensus genome sequence where the heterozygous sites were represented with IUPAC ambiguity codes was generated using bcftools v1.17 (Danecek et al. 2021). The consensus sequence was converted to the input format of PSMC by applying a bin size of 50 bp.

For PSMC, we included data from all chromosome-size scaffolds except for the X chromosome (scaffold 4). PSMC was run using the parameter settings *-N25 -t15 -r5 -p “4+25*2+4+6”*. We also explored including more parameters (*-p “4+45*2+4+6”* and *-p “4+65*2+4+6”*) as well as splitting the first parameter (e.g., *-p “2+2+25*2+4+6”*), as recommended by (Hilgers et al. 2024). None of these settings changed the general results, so we chose to use the default parameters.

Finally, we calibrated PSMC results using a generation time of 1.5 years and a mutation rate of 5.7*10^-9^ per generation. This mutation rate is based on the assumption that garden dormice exhibit a similar mutation rate as mice, for which the rate is well established (Uchimura et al. 2015; Milholland et al. 2017). The mean generation time (1.57 years) was calculated based on a yearly reproductive cycle starting at age one, a maximum age of 5 years, and an average yearly survival rate of 0.38 (Schaub and Vaterlaus-Schlegel 2001).

### Population genomics

#### Genotyping, Filtering and Variant Calling

Full methods for processing the raw GBS data, including trimming and alignment, are described in the Supplementary Methods. Variant calling was performed with the ref_map.pl pipeline for reference aligned reads in Stacks v2.65 (Rochette et al. 2019) with additional filtering parameters given for the populations program: minor allele frequency of 5%, a minimum percentage of individuals per population of 80, and writing one random SNP per locus to remove strong physical linkage disequilibrium (-X “populations: -r 0.80 --min-maf 0.05 --write-random-snp”). Additional variant filtering was performed with VCFtools v0.1.16 (Danecek et al. 2011) to remove indels and the X Chromosome and only keep biallelic SNPs with a depth per individual between 6 and 50 (flags: ‘--remove-indels --min-alleles 2 --max-alleles 2 --min-meanDP 6 --max-meanDP 50 --minDP 6 -- maxDP 50’). After filtering, we removed samples with > 80% missing data to generate our final VCF file.

Past mitochondrial DNA work has classified four distinct garden dormouse clades within its European range (Iberian, Italian, Western European, and Alpine; (Perez et al. 2013). We followed these clade designations in assigning *a priori* population identity but expanded the ‘Western European’ category by delineating individuals within this clade as Northwest, the Harz Mountains, Northeast, Central, and Russia (Fig. 4A) as aligning to known garden dormouse distribution within this region (Meinig and Büchner 2012).

Because linkage disequilibrium (LD) can influence some analyses such as principal components-based analyses (PCAs), we created a pruned SNP set using the +prune plugin in BCFtools (Danecek et al. 2021) using an LD threshold of r^2^ < 0.5 and a sliding window of 1000, which was applied to analyses that rely on allele frequencies for population inferences. We hereafter refer to the ‘full’ and ‘pruned’ SNP sets.

Excessive numbers of related individuals can bias some population genomics analyses ((O’Connell et al. 2019); (Wang 2018)). Because our sample collection was not always random, we anticipated that several individuals within our sample set were related, and therefore estimated pairwise relatedness within sampling regions with the full SNP set pruned by r^2^ < 0.2 using the KING-robust method ((Manichaikul et al. 2010) in SNPRelate. We then removed 1 from each pair with a kinship estimate > 0.35, which corresponds to a first-order familial relationship (sibling pair, parent-offspring). This sample set was used for all subsequent analyses.

#### Genetic Diversity

Genetic diversity estimates can be biased by a number of factors, including allele frequencies, unbalanced sample size, and missing data (Schmidt et al. 2021). We calculated observed (H_obs_) and expected (H_exp_) heterozygosity, nucleotide diversity (**π**), number of polymorphic loci, and the inbreeding coefficient F_IS_ with the pruned SNP set using the ‘population’ function in stacks (Rochette et al. 2019). Because biases due to unbalanced sample sizes can confound comparisons among populations, we additionally report H_exp_ derived from genotype-wide autosomal data, which is not subject to the same bias (Schmidt et al. 2021; Sopniewski and Catullo 2024) Autosomal heterozygosity was generated from the aligned Stacks catalog using the ‘population’ function with a minor allele count of 2 to omit singletons and with one random SNP selected per locus. To further minimize the effects of differences in sample size, we calculated SNP-based genetic diversity parameters for the Northwest population in four separate runs of *n* = 12 samples per run and report the mean values. We investigated genetic differentiation by calculating pairwise values of Weir and Cockerhams’ unbiased estimator of *F*_ST_ (Weir and Cockerham 1984) in the R package hierfstat (Goudet and Goudet 2014) using the pruned SNP set and with 95% confidence intervals generated with 1,000 bootstrap iterations.

#### Population Structure

We investigated phylogenetic distance among samples and groups using the full SNP set by inferring a phylogeny in RAxML version 8.2.12 (Stamatakis 2014) using the ASC-GTRCAT model with Lewis correction for ascertainment bias with 100 bootstrap replicates. We plotted the tree in the R 4.3 (R Core Team 2018) package GGTREE (Yu et al. 2017) with a customized script formulated for SNP marker data (Severn-Ellis et al. 2020).

To explore population structure, we first used the pruned SNP set to visualize population clustering via PCA in the R package *adegenet* (Jombart and Bateman 2008)). We estimated the most probable number of genetic clusters K within our sample set using the full SNP set. First, we implemented a discriminant analysis of principal components (DAPC) using the ‘find.clusters’ function in *adegenet* ((Jombart and Bateman 2008)) to estimate optimal number of *K* testing *K* = 1–8. The most probable number of clusters was identified as the one with the lowest Bayesian Information Criterion (BIC) value. To minimize overfitting, we first determined the optimal number of principal components (PCs) using cross-validation and then ran DAPC retaining the optimal number of PCs and using the identified optimal number of genetic clusters as *K* to predict group assignment for each sample without *a priori* population information. Five principal components (PCs) were retained to explain approximately 52% of the total retained variation without our dataset.

We also used STRUCTURE v2.3.4 (Pritchard 2000) within the program *Structure_threader* (Pina-Martins et al. 2017) to estimate the value of *K*. We ran STRUCTURE with the admixture model and no prior population information, with an optimal burn-in of 10^4^ steps and 10^4^ additional steps, and for values of *K* from 1 to 8 with 10 replicates per iteration of *K*. We used the R package pophelper (Francis 2017) to align runs, plot output, and determine the most likely value of *K* via both ΔK ((Evanno et al. 2005) and mean probability (MeanLn*P(K)*). STRUCTURE is known to be sensitive to differences in sample size in populations ((Puechmaille 2016), which was a factor in our dataset. To eliminate the potential effects of sampling bias on our results, we also ran STRUCTURE using the same conditions for a random subset of *n* = 5–6 individuals per population, omitting the Russian and Italian samples from the dataset, for values of *K* from 1 to 5.

#### Recent Demographic History Inferred from the Site-frequency Spectrum

We inferred demographic history among our individual populations using a sequential Markovian coalescent approximation approach in *fastSimcoal2* v2.08 (Excoffier and Foll 2011; Excoffier et al. 2021). We omitted the Central and Iberian Peninsula populations and the Italian and Russian samples due to their low sample sizes. To generate the input site frequency spectrum (SFS) files, we used the distribution of minor allele frequencies in the full SNP set to create a folded SFS for each population with the R program dartR 2.9.7 (Gruber et al. 2018), using nearest-neighbor imputation to replace missing data. While *fastSimcoal2* can be used to infer complex demographic co-histories, we opted to test models for each population separately to avoid model overparameterization, given our small sample sizes (Alps, n = 9; Harz, n = 12; Northeast, n = 7; Northwest, n = 48). For each population, we compared three models: “null”, “instantaneous size change” (a.k.a., bottleneck), and “incremental size change”. The null model assumed a constant population size over time and inferred only one parameter, contemporary effective population size (N_e_). The instantaneous size change model assumed a rapid population decline and inferred four parameters: ancestral N_e_, time of the bottleneck (in generations), population size during the bottleneck, and contemporary N_e_. The incremental size change model assumed either steady population growth or decline over time and inferred five parameters: ancestral N_e_, time of ancestral N_e_ (in generations), population growth rate (with a negative value indicating a positive growth rate), rate of change of N_e_, and contemporary N_e_. For all models, we assumed a generation time of 1.5 years and a mutation rate of 5.7×10^-9^ . Parameter ranges were uniformly distributed between 10 and 10^5^ We ran 100 independent iterations of each model per population with 70 EMC cycles and 1×10^6^ simulations per model run. We selected the best run within each model as the one with the highest likelihood (smallest difference between maximum likelihood and obtained likelihood) using the custom script fsc-selectbestrun.sh (https://raw.githubusercontent.com/speciationgenomics/scripts/master/fsc-selectbestrun.sh). Because maximum likelihood can be influenced by parameter size, model selection within each population was then performed using Akaike Information Criterion (AIC) comparisons, which account for the number of parameters in the selection process, with AIC calculated using the custom script calculateAIC.sh (https://github.com/speciationgenomics/scripts/blob/master/calculateAIC.sh).

## Data Access

Genomic and transcriptomics read data, our annotated genome and the trimmed ddRAD reads have been submitted to the NCBI BioProject database (https://www.ncbi.nlm.nih.gov/bioproject/) under accession number PRJNA1142780. The haplotype assemblies, gene and transposable element annotations are available for download at https://genome.senckenberg.de/download/GardenDormouse/. TOGA annotations are available at https://genome.senckenberg.de/download/TOGA/. We also provide a genome browser showing our genomes and all annotations at https://genome.senckenberg.de/.

## Code Access

Most of our analyses relied on publicly available tools and methods. New scripts are available at https://github.com/pabyerly/TBG_GardenDormouse and as Supplementary Code.

## Acknowledgment

This work was supported by the LOEWE-Centre for Translational Biodiversity Genomics (TBG) funded by the Hessen State Ministry of Higher Education, Research and the Arts (LOEWE/1/10/519/03/03.001(0014)/52). Funding was also provided by the German Federal Agency for Nature Conservation (BfN) with resources from the Federal Ministry for the Environment, Nature Conservation, Nuclear Safety and Consumer Protection (BMUV), as part of a German-wide project led by the Bund für Umwelt und Naturschutz BUND (Friends of the Earth Germany), Justus Liebig University Gießen, and the Senckenberg Gesellschaft für Naturforschung in the Federal Programme for Biological Diversity called ‘In Search of the Garden Dormouse’.

We are very grateful to many people who have helped with sample collection: Peter Adamik, Hermann Ansorge, Jonas Astrin, Simon Capt, Stefanie Jessolat, Jeroen van der Kooij, Ines Leonhard, Gernot Segelbacher, David Stille, Adria Viñals Domingo, Frank Zachos, Alain Frantz, Olga Grigoryeva, Maurice La Haye, Roel Baets, Kasper Van Acker, Caren Raditz, Lutz Nielen, Stefanie Erhardt, Jürg Paul Müller, Andrea Krug, Danilo Hartung, Christine Thiel-Bender, Ferry Böhme, Julia Hofmann, Sarah Beer, Jörg und Elli Hacker, Otfried Wüstemann, Sonja Klein, Thomas Doebel, Martina Klotz, Hartmut Schmid, the Schwarzwald National Park, the Gran Paradiso National Park, the Museo di Scienze Naturali dell’Alto Adige and Museo di Storia Naturale La Specola di Firenze, Antoni Arrizabalaga, Natural Sciences Museum of Granollers, as well as numerous students and citizen scientists without whom this work would have not been possible.

## Author contributions

A.T., C.N., M.H. designed and supervised the study, labwork and data analysis. C.G., A.B.H., C.G., C.B, and H.J.B. performed labwork. P.A.B., E.L., L.H., S.L., S.W., and T.S. conducted statistical and bioinformatic analysis. S.B., J.L., H.M., E.M.F-P., S.P.S., A.M., S.B., G.V., T.B., L.F., L.V., and S.A.M. contributed samples, supervised sample collection, and/or provided expert species knowledge. P.A.B., A.T., E.L., L.H., and M.H. wrote the manuscript with input from all authors.

