## Supplementary Figures and Methods for "Haplotype-resolved genome and population genomics of the threatened garden dormouse in Europe"

### Supplemental Material for
### Haplotype-resolved genome and population genomics of the threatened garden dormouse in Europe


Paige A. Byerly[1,2#*], Alina von Thaden[1,2#], Evgeny Leushkin[1,3], Leon Hilgers[1,3], Shenglin Liu[1,3], Sven Winter[4,5], Tilman Schell[1,3], Charlotte Gerheim[1,3], Alexander Ben Hamadou[1,3], Carola Greve[1,3], Christian Betz[6], Hanno J. Bolz[6], Sven Büchner[8], Johannes Lang[8], Holger Meinig[8], Eva Marie Famira-Parcsetich[8], Sarah P. Stubbe[8], Alice Mouton[9], Sandro Bertolino[10], Goedele Verbeylen[11], Thomas Briner[12], Lídia Freixas[13], Lorenzo Vinciguerra[14], Sarah A. Mueller[15], Carsten Nowak[1,2], and Michael Hiller[1,3,7*]

[1] LOEWE Centre for Translational Biodiversity Genomics, Senckenberganlage 25, 60325 Frankfurt, Germany

[2] Conservation Genetics Group, Senckenberg Research Institute and Natural History Museum Frankfurt, Clamecystraße 12, 63571 Gelnhausen, Germany

[3] Senckenberg Research Institute, Senckenberganlage 25, 60325 Frankfurt, Germany

[4] Senckenberg Biodiversity and Climate Research Centre, Senckenberganlage 25, 60325 Frankfurt am Main, Germany

[5] Research Institute of Wildlife Ecology, University of Veterinary Medicine Vienna, Savoyenstr. 1, 1160 Vienna, Austria

[6] Bioscientia Human Genetics, Institute for Medical Diagnostics GmbH, Ingelheim, Germany

[7] Institute of Cell Biology and Neuroscience, Faculty of Biosciences, Goethe University Frankfurt, Max-von-Laue-Str. 9, 60438 Frankfurt, Germany

[8] Justus-Liebig-University Giessen, Clinic for Birds, Reptiles, Amphibians and Fish, Working Group for Wildlife Research, Frankfurter Strasse 114, 35392 Giessen, Germany

[9] Socio-économie, Environnement et Développement (SEED), University of Liege (Arlon Campus Environment), Belgium

[10] Department of Life Sciences and Systems Biology, University of Turin, 10123 Torino, Italy

[11] Natuurpunt Studie vzw, Mammal Working Group, Michiel Coxiestraat 11, 2800 Mechelen, Belgium

[12] Naturmuseum Solothurn, Klosterplatz 2, 4500 Solothurn, Switzerland

[13] BiBio Research Group, Natural Sciences Museum of Granollers, C/ Francesc Macià 51, 08402 Granollers, Catalonia, Spain.

[14] Naturmuseum St.Gallen, Rorschacher Strasse 263, 9016 St. Gallen, Switzerland

[15] Division of Evolutionary Biology, Faculty of Biology, LMU Munich, Germany

### contributed equally to the manuscript

The Supplemental Material contains

- Figures S1 to S7
- Supplemental Methods

Tables S1 and S2 are provided as sheets in an Excel file.

**Table legends**

Table S1: Assembly statistics

The table lists Sciuromorpha species for which genome assemblies are available on NCBI or DNAzoo, together with their contig/scaffold N50 values and gene completeness benchmarks from TOGA and compleasm. These assemblies have been included in the comparison in Figure 2. Compleasm distinguishes between two types of fragmented genes: F = only one part is present; I = the whole BUSCO gene is present but in different parts in the assembly.

Table S2: Sample metadata.

Metadata for analyzed samples, including the sample identification number (ID), date the sample was collected (SampleDate), location where the sample was collected (Country, Region, latitude, longitude), the sample provider (collector), assigned population for genomic analysis (visual representation in Figure 4A), frequency of missing SNPs (%miss), and per sample mean read depth (depth). NA in %miss and depth indicates that sample was not used in analysis.

**Figure S1: Sequencing coverage, observed heterozygosity and repetitive sequence content.**

The three plots show the sequencing coverage (top), observed heterozygosity (middle) and repetitive sequence content (bottom) using a sliding window size of 1 Mb across the chromosome assemblies of the haplotype 1 (A) and haplotype 2 (B). Chromosomes are ordered independently according to their size in haplotype 1 (A) and haplotype 2 (B). In (B), we list the names of the homologous haplotype 1 chromosomes in parentheses.

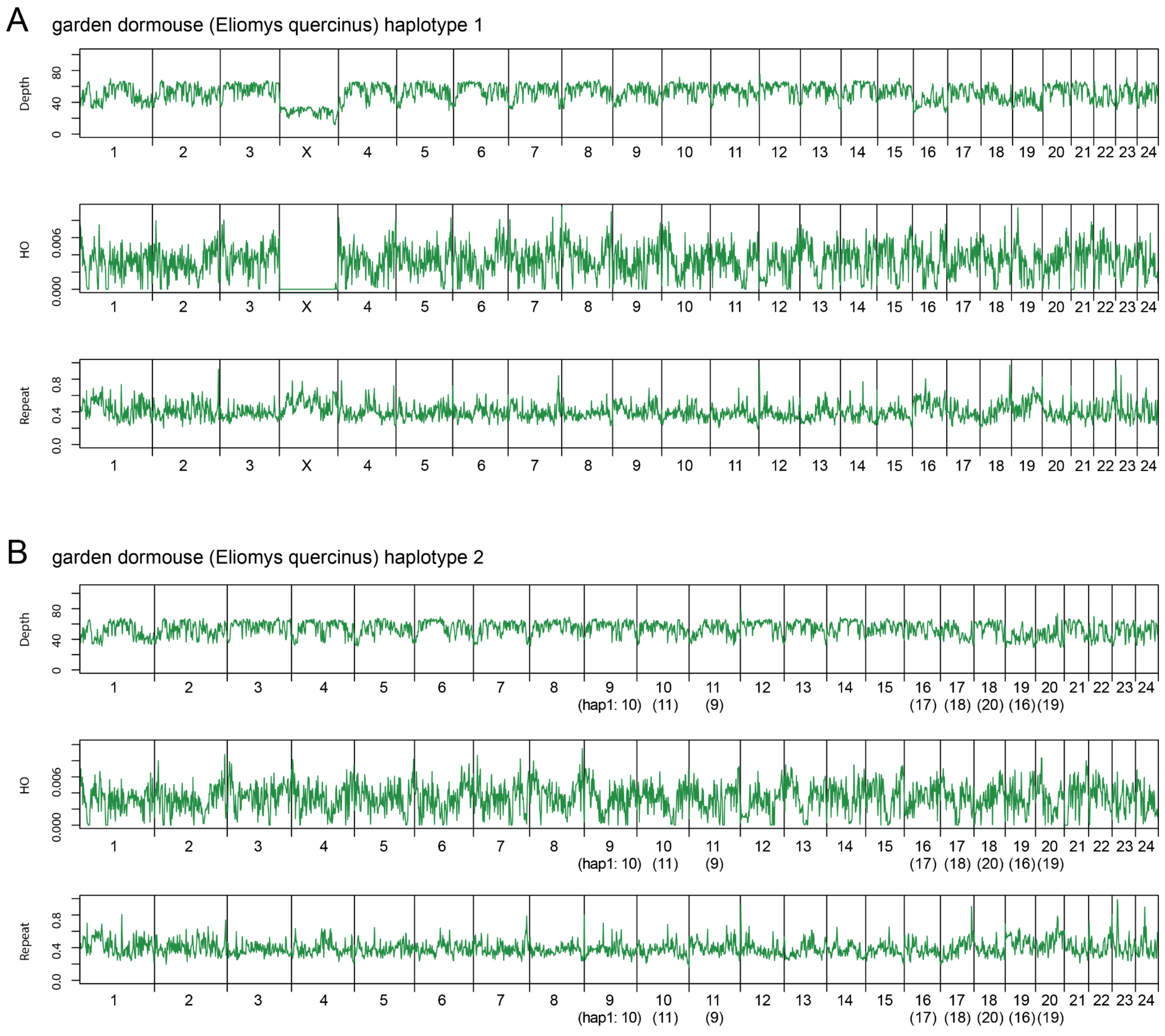

**Figure S2: Maximum likelihood phylogeny of the garden dormouse in Europe based on 41,775 nuclear SNP loci as inferred in RAxML.**
Colors within nodes represent geographic sampling regions.

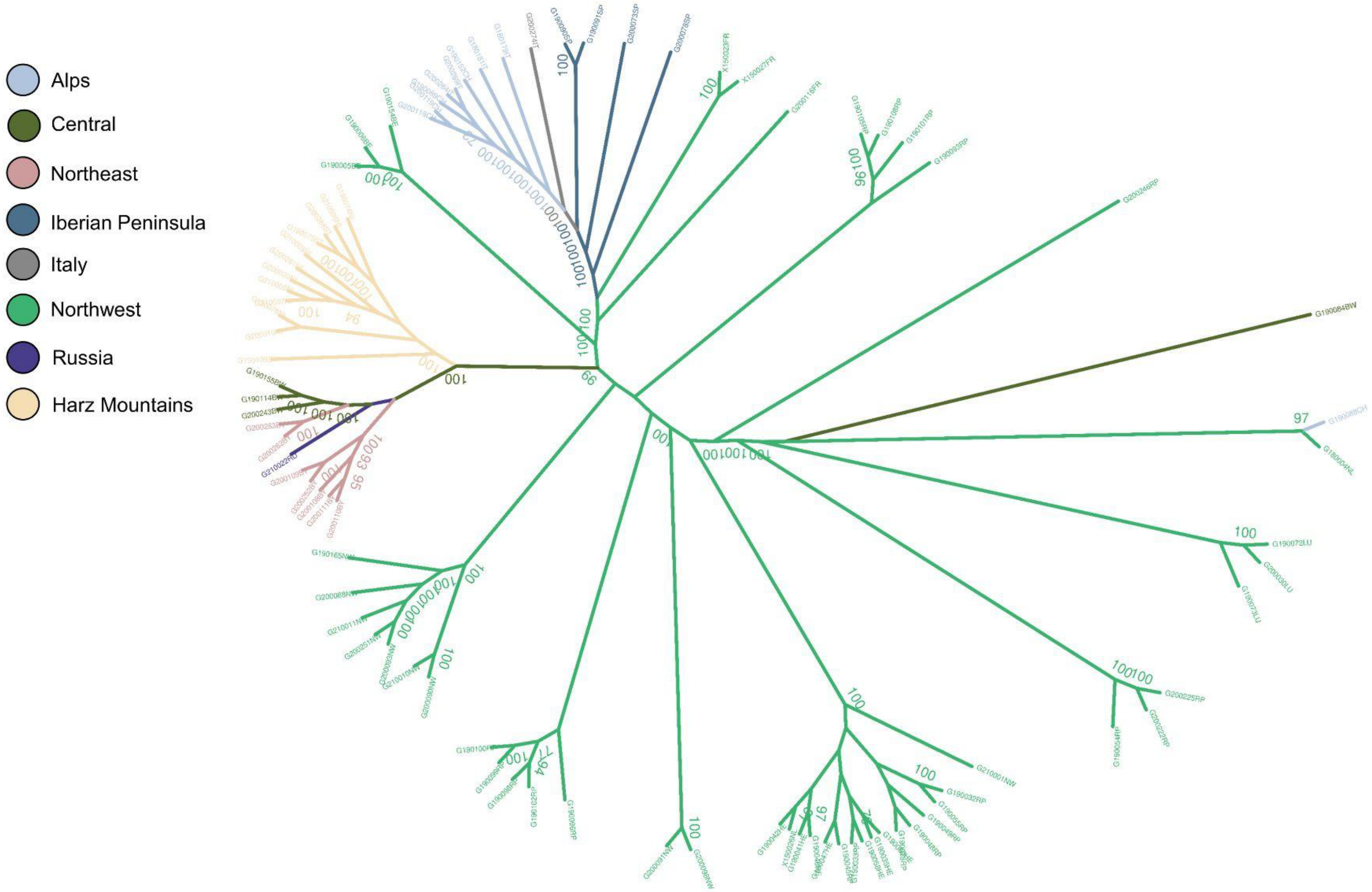

**Figure S3. Discriminant analysis of principal components (DAPC).**
(A) BIC plotted to identify the most likely value of genetic clusters *K*, which is *K* = 4.
(B) Garden dormouse genomic data plotted as *K* = 4 on two discriminant functions, with shapes representing individuals, colors designating predicted grouping, and the x- and y-axes describing the first two discriminant functions.
(C) Predicted group membership for groups 1–4.

**A.**

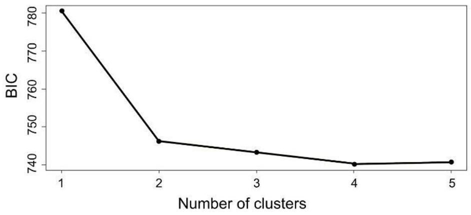


**B.**

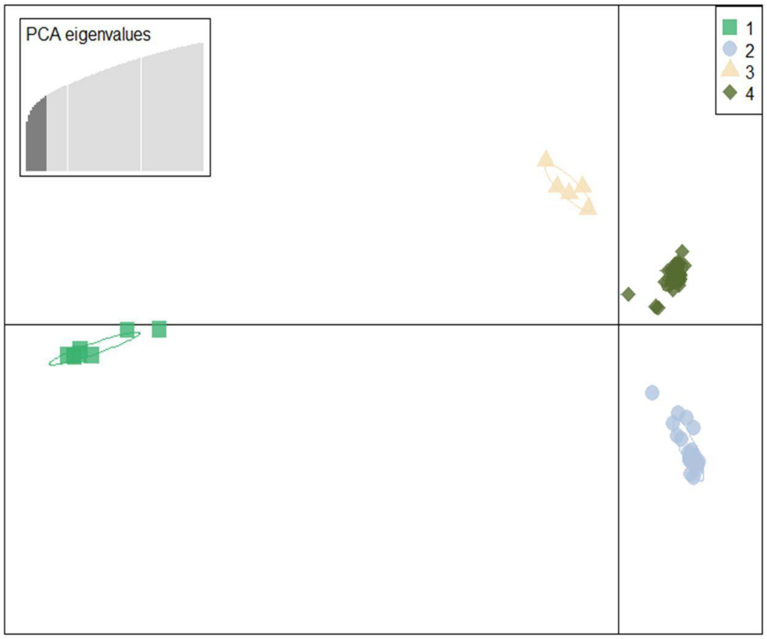


**C.**

| | 1 | 2 | 3 | 4 |
|---|---|---|---|---|
| **Northwest** | 0 | 0 | 48 | 0 |
| **Alps** | 0 | 8 | 1 | 0 |
| **Harz Mountains** | 12 | 0 | 0 | 0 |
| **Central** | 3 | 0 | 1 | 0 |
| **Iberian Peninsula** | 0 | 0 | 0 | 4 |
| **Northeast** | 7 | 0 | 0 | 0 |
| **Italy** | 0 | 0 | 0 | 1 |
| **Russia** | 1 | 0 | 0 | 0 |

**Figure S4. Output of R package pophelper showing the Evanno method for estimating likeliest value of *K* from the STRUCTURE output.**
(A-D) represent four different *K*-value selection metrics for hypothesized values of *K* ranging from 1 to 8.

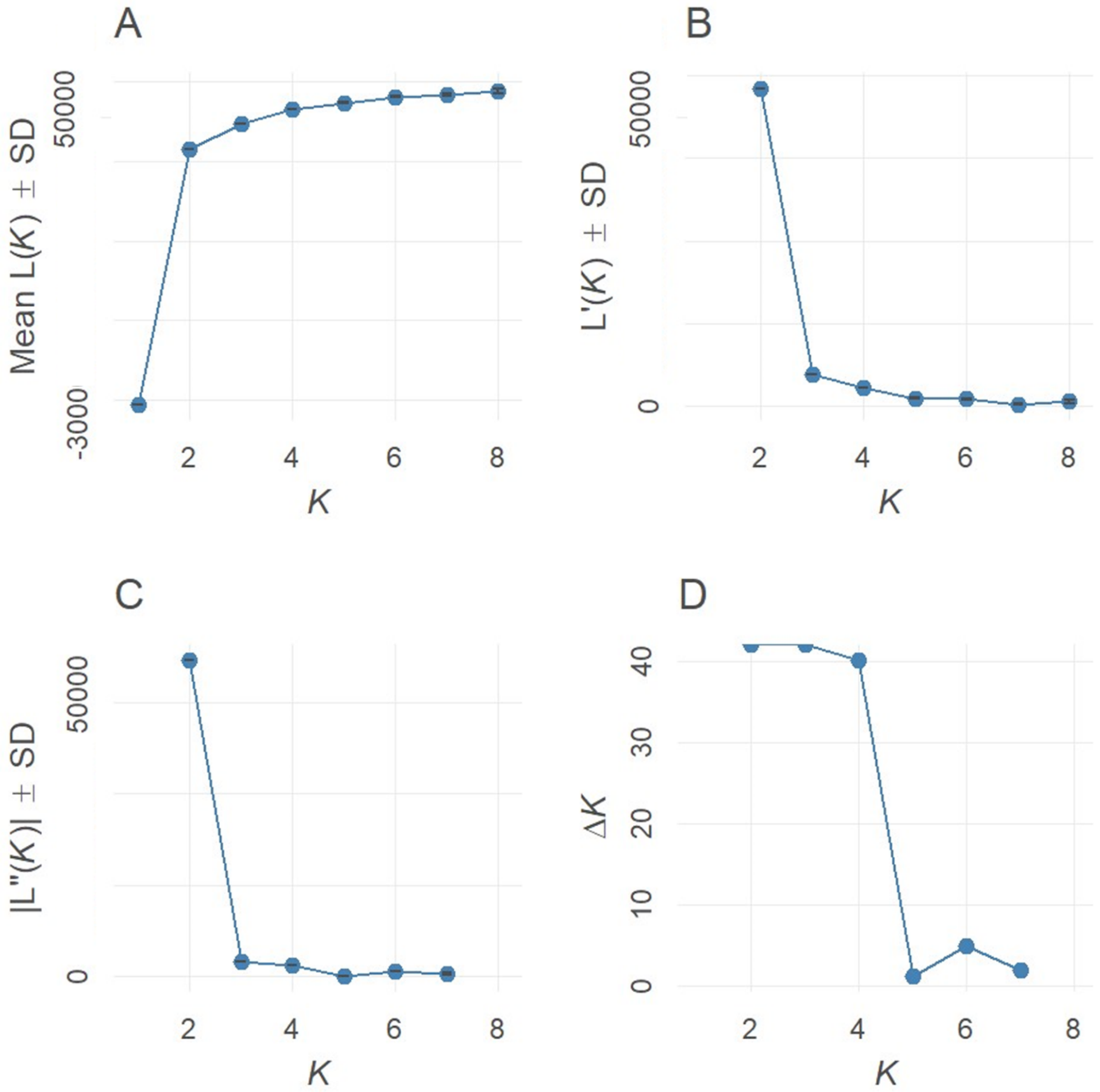

**Figure S5. Genetic clustering inferred by Bayesian structure analysis for *K* ranging from 2 to 8 in STRUCTURE.**

The x-axis represents geographic sampling location and the y-axis the proportion of group membership.

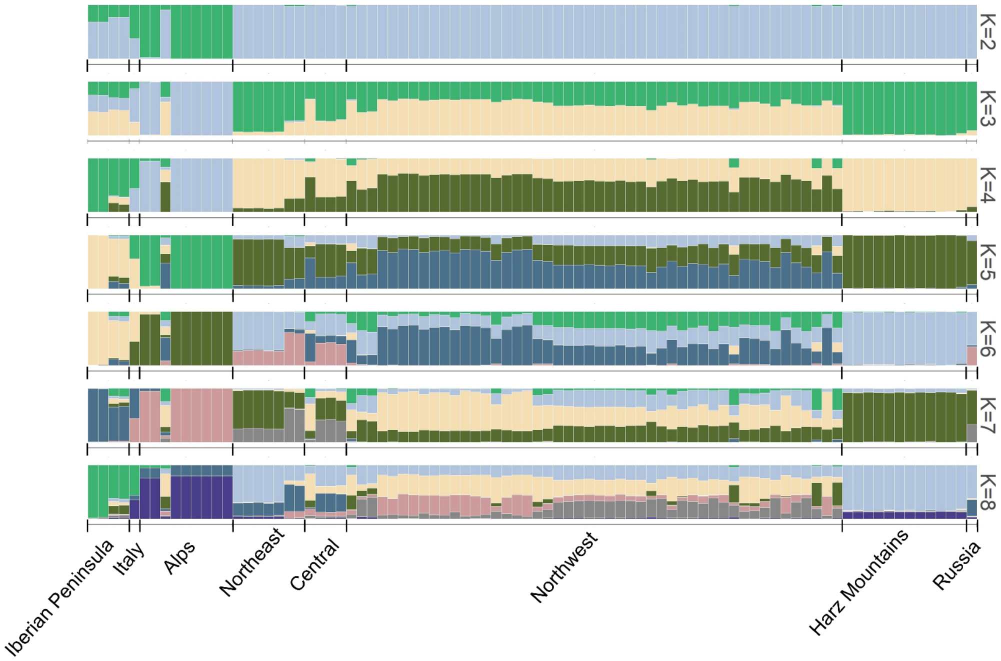

**Figure S6. Genetic clustering of the subset dataset (*n* = 4–5 per population) inferred by Bayesian structure analysis for *K* ranging from 2 to 5 in STRUCTURE.**
The x-axis represents the geographic sampling location and the y-axis the proportion of group membership.

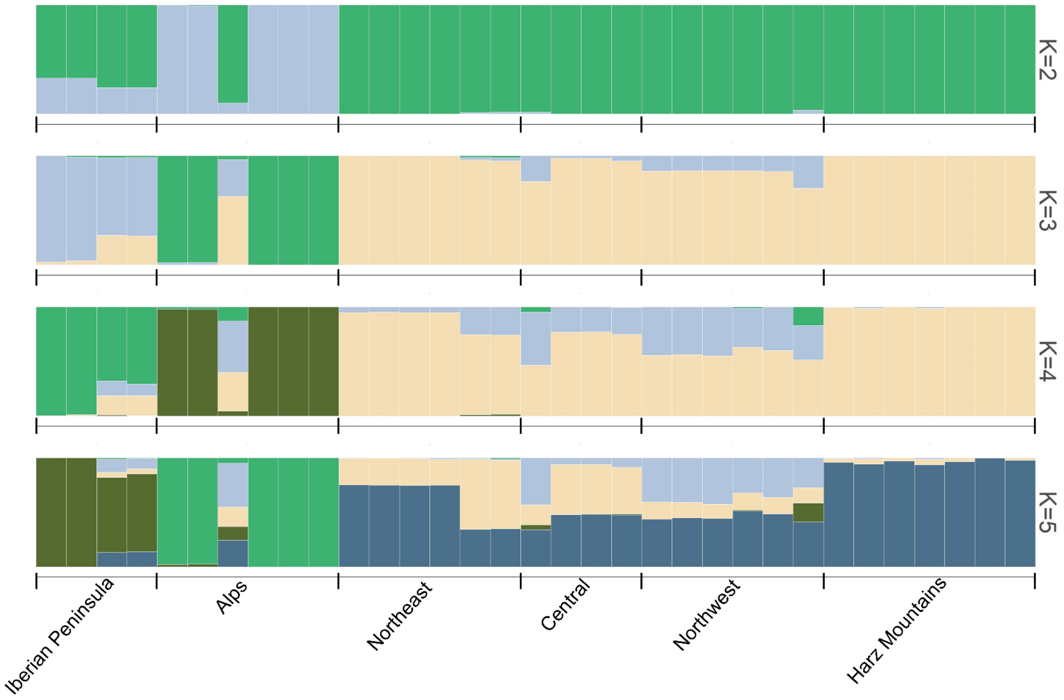

**Figure S7. Graphical representation of coalescent models compared to infer the most likely demographic histories for four garden dormouse populations in fastsimcoal2.**

Models tested were: **a.** null model depicting a constant population size over time with one inferred parameter, contemporary effective population size (**$N_e$CON**); **b.** instantaneous size change model assuming a rapid population decline with potential for **b1.** population recovery (bottleneck) or **b2.** no population recovery, and with four inferred parameters: ancestral $N_e$ (**$N_e$ANC**), time period of the bottleneck in generations (**TBOT**), $N_e$ during the bottleneck (**$N_e$BOT**), and **$N_e$CON**; and **c.** incremental size change model assuming either **c1.** steady population growth or **c2.** decline over time, and inferring five parameters, including: **$N_e$ANC**, time period of ancestral $N_e$ in generations (**TBOT**), population growth rate and rate of change (**r** and **ROC)**, and **$N_e$CON**. For all models, we assumed a generation time of 1.5 years and a mutation rate of $5.7*10^{-9}$ .

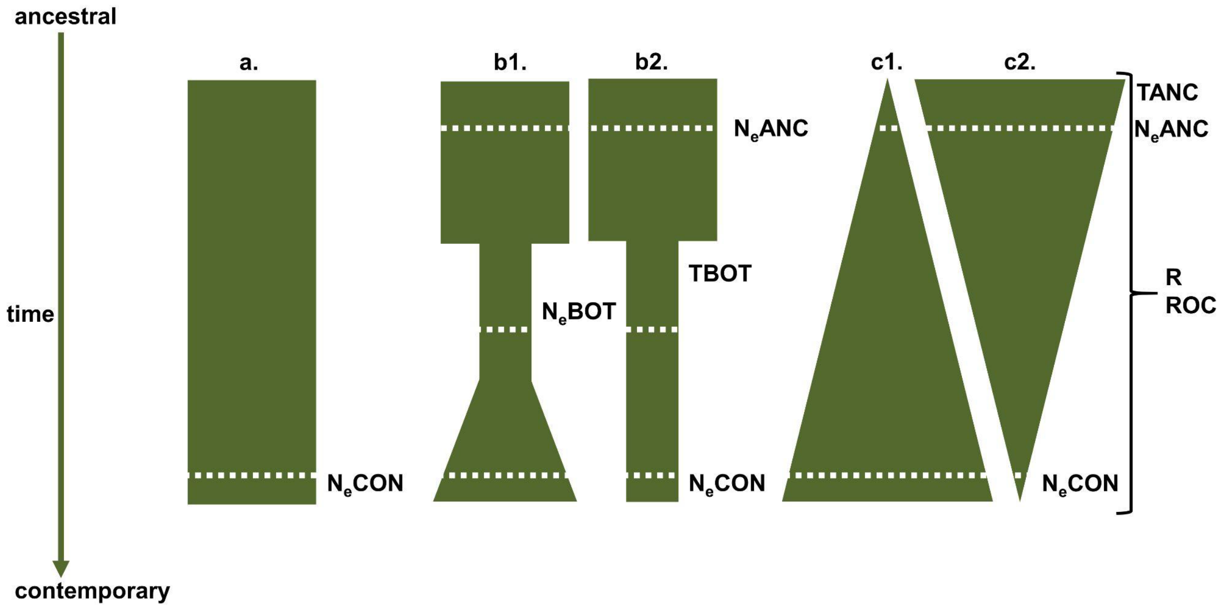

#### Supplemental Methods

##### Genome sequencing

High molecular weight genomic DNA was extracted from heart tissue according to the protocol of (Sambrook and Russell 2001) for the first library and from lung tissue using the PacBio Nanobind Tissue Kit (PacBio, Menlo Park, CA, USA) for the second library. DNA concentration and DNA fragment length were assessed using the Qubit dsDNA BR Assay kit on the Qubit Fluorometer (Thermo Fisher Scientific) and the Genomic DNA Screen Tape on the Agilent 4150 TapeStation system (Agilent Technologies), respectively. A SMRTbell library was prepared for each tissue according to the instructions of the SMRTbell Express Prep Kit v3.0. Total input DNA was approximately 4 µg per library.

Quality control of the final libraries was performed on an Agilent Femto Pulse system (Agilent, Waldbronn Germany). Annealing of sequencing primers, binding of sequencing polymerase, and purification of polymerase-bound SMRTbell complexes were performed using the Revio polymerase kit (PacBio, Menlo Park, CA, USA). Loading concentration for sequencing was 225 pM. Long-read whole-genome sequencing was performed on a PacBio Revio instrument (PacBio, Menlo Park, CA, USA), 24 h HiFi sequencing, 15 kb median read length, Q30 median read quality. Two PacBio Revio SMRT cells generated in total 124.4 Gb in HiFi reads that were used for contig genome assembly.

To generate long-range data for scaffolding, we used the Arima High Coverage Hi-C Kit v01 (Arima Genomics) according to the Animal Tissue User Guide for proximity ligation using approximately 110 mg of heart tissue from the same individual. The proximally-ligated DNA was then converted into an Arima High Coverage Hi-C library according to the protocol of the Swift Biosciences® Accel-NGS® 2S Plus DNA Library Kit. The fragment size distribution and concentration of the Arima High Coverage Hi-C library was assessed using the TapeStation 4150 (Agilent Technologies) and the Qubit Fluorometer and Qubit dsDNA HS reagents Assay kit (Thermo Fisher Scientific, Waltham, MA), respectively. The library was sequenced on the NovaSeq 6000 platform at Novogene (UK) using a 150 paired-end sequencing strategy, resulting in an output of 117 Gb.

##### Transcriptome sequencing

Total RNA was isolated from the eight organs using TRIzol reagent (Invitrogen) according to the manufacturer's instructions. The quality and concentration of each extraction was assessed using the TapeStation 4150 (Agilent Technologies) and the Qubit Fluorometer with the RNA BR Reagents Assay Kit (Thermo Fisher Scientific, Waltham, MA). The RNA extractions were then sent to Novogene (UK) for Illumina paired-end 150 bp RNA-seq of a cDNA library (insert size: 350 bp) with a respective expected output of 9 Gb.

For the preparation of Iso-Seq libraries using the SMRTbell Prep Kit 3.0, only RNA extractions with an RNA integrity number (RIN) > 7 were used, as recommended by the manufacturer's protocol. Two libraries were prepared by pooling RNA from kidney, liver, lung or bladder, heart and testis at approximately 90 ng each, with the exception of lung RNA at approximately 60 ng, resulting in two pooled libraries. These two Iso-Seq libraries were then each loaded onto a SMRT cell sequenced in CCS mode using the Sequel System IIe with the Sequel II Binding Kit 3.1 (Pacific Biosciences, Menlo Park, CA) at the Genome Technology Center (RGTC) at Radboudumc (Nijmegen, The Netherlands). The libraries were loaded to an on-plate concentration of 80 pM each by diffusion loading.

**Population genomics: Genotyping, Filtering and Variant Calling**

Raw GBS data were trimmed using fastp v0.23.4 (Chen et al. 2018) with enabled base correction and low complexity filter to remove sequencing adaptors and poly(G) stretches at the end of reads. A 4 bp sliding window was employed to detect regions of poor quality (Phred score < 15). We removed reads if they fit into one of the following categories: read length below 36 bp, reads with > 40% low-quality bases, and reads with 5 or more undetermined bases (Ns). The trimmed reads were then mapped against haplotype 1 of our assembly using BWA-MEM v0.7.17-r1188 (https://github.com/lh3/bwa). Mate coordinates in the resulting mapping files were filled in, the files were sorted by position, and indexed using SAMtools v.1.18 (Danecek et al. 2021).
